# A Dynamic Oligomerization Network Coordinates Hemagglutinin-Mediated Membrane Fusion on Influenza Virions

**DOI:** 10.64898/2026.03.23.713829

**Authors:** Yong Chen, Zheyuan Zhang, Zirui Zhao, Hanguang Liu, Haifan Zhao, Rui Liang, Cheng Peng, Jialu Xu, Yutong Song, Xu Tan, Sai Li

## Abstract

Influenza A virus (IAV) entry is mediated by the trimeric glycoprotein hemagglutinin (HA), which undergoes low-pH-triggered conformational rearrangements to drive membrane fusion. Although biochemical and biophysical studies have long suggested that HA trimers function cooperatively, the structural basis for this cooperativity on intact virions has remained unclear. Here, using cryo-electron tomography (cryo-ET) and subtomogram averaging (STA), we determine the in situ structure of prefusion HA on native virions at 3.58 Å resolution, enabling atomic modeling of membrane-anchored HA. We show that neighbouring HA trimers engage in lateral interactions mediated by HA1 subunits, assembling into flexible dimeric, pentameric, and hexameric organizations that form locally ordered lattices on the viral surface. A 6.0 Å reconstruction of the HA-dimer reveals the molecular interfaces underlying these assemblies. Disruption of the dominant HA1-HA1 interaction markedly impairs viral entry and slows membrane fusion kinetics. Together, these findings define a virion-level mechanism for coordinated HA activation and establish a structural framework for understanding cooperative membrane fusion by class-I viral fusion proteins.

## Introduction

Enveloped viruses deliver their genomes into host cells through membrane fusion, a process mediated by three major classes of viral fusion proteins^1^. Class-I fusion proteins, encoded by viruses from the *Orthomyxoviridae*, *Paramyxoviridae*, *Coronaviridae*, and *Retroviridae* families, are predominantly α-helical homotrimers. Class-II fusion proteins, characteristic of *Togaviridae*, *Phleboviridae*, *Flaviviridae*, and *Nairoviridae*, exhibit multiple oligomeric states and are primarily composed of ß-sheets. Class-III fusion proteins, found in *Orthoherpesviridae* and *Rhabdoviridae*, are homotrimers containing both α-helical and ß-sheet elements. Given the significant energy required to bend the lipid bilyaers, drive hemifusion, and ultimately open a fusion pore, the coordinated action of multiple fusion proteins is expected. In class-II systems, such cooperativity can be pre-organized in the prefusion state, as they oligomerize into higher-order complexes on the viral envelope^2^. In contrast, prefusion class-I and class-III fusogens were conventionally thought to distribute randomly. However, accumulating evidence suggests that multiple class-I trimers are required to drive efficient membrane fusion^3–7^, raising questions on how spatially dispersed trimers are recruited and coordinated at fusion sites.

Recent structural studies revealed oligomeric assemblies of prefusion class-I trimers on intact enveloped viruses. For example, prototype foamy virus (PFV) Envs form pentameric and hexameric assemblies of trimers^8^; respiratory syncytial virus (RSV) fusion (F) proteins adopt dimer-of-trimers configurations^9^; influenza C virus (ICV) hemagglutinin-esterase-fusion (HEF) proteins assemble into hexagonal lattices^10^, and IAV HAs form paired arrangements^11^. Similar oligomeric organizations have also been described for class-III trimers, including herpes simplex virus (HSV-1) glycoprotein B (gB)^12^ and vesicular stomatitis virus (VSV) glycoprotein (G)^13^, which form pentameric and hexameric assemblies. These observations suggest that lateral interactions among prefusion fusion proteins may represent a conserved organizational principle across enveloped viruses.

IAV is an enveloped, negative-sense RNA virus from the *Orthomyxoviridae* family. The viral envelope is decorated with two major glycoproteins, HA and neuraminidase (NA). Beneath the membrane, matrix protein 1 (M1) polymerizes into a layer tightly associated with the viral envelope^14^, where it interacts with viral ribonucleoproteins (RNPs)^15^ and the cytoplasmic tails of HA and NA^16^. Extensive structural and functional insights into IAV glycoproteins have primarily been derived from studies of soluble ectodomains^17,18^. HA comprises the receptor-binding HA1 subunit and the membrane fusion HA2 subunit, a class-I viral fusogen. Following endocytosis, exposure to the acidic endosomal environment triggers HA refolding, resulting in membrane fusion^18^. NA, a tetrameric sialidase, cleves sialic acids to balance HA-mediated attachment and facilitate viral mobility and release^19^.

Mechanistic studies have demonstrated that multiple HA trimers cooperate during membrane fusion. Cryo-ET analyses of IAV virus-liposome fusion intermediates have characterized the sequence of membrane remodeling^20,21^, and revealed fusion pores encircled by radially arranged HAs^7^. Single-molecule experiments suggest that approximately three to six HAs are required for productive membrane fusion^4,5,22,23^. Low-resolution on-virion reconstruction of HA-dimers suggested they are formed by lateral interactions between the HA1 subunits^11^. However, owing to limited structural resolution and the absence of functional validations, the molecular determinants, structural mechanisms, and functional consequences of HA oligomerization on intact IAV surface remain incompletely understood.

Here, we applied cryo-ET to determine the on-virion structures of HA from influenza virus A/Puerto Rico/8/34 (H1N1, PR8) at near-atomic resolution. Our reconstructions resolve the membrane-proximal stalk region and enabled atomic modeling of the full HA ectodomain in situ, revealing conformational features distinct from previously characterized soluble HA structures. We further identify multiple modes of HA oligomerization on the viral surface, including dimer-, pentamer-, and hexamer-of-trimers assemblies, and determine their structures at 6.0 Å, 8.0 Å, and 9.7 Å resolution, respectively. Structural analysis reveals two HA1-HA1 interaction interfaces that underpin oligomer formation, and mutational studies demonstrate that these interactions play critical roles in viral entry and membrane fusion. Collectively, our findings define a dynamic structural landscape of glycoproteins on the IAV envelope and establish a mechanistic framework for HA-mediated cooperative fusion.

## Results

### HA structures on cell-derived virions

PR8 virions were propagated in MDCK cells, isolated in neutral-pH buffer, and imaged by cryo-ET (Figure 1A). From approximately 85,000 HA particles extracted from 446 virions, the prefusion HA ectodomain structure was resolved to an overall resolution of 4.09 Å (Figures 1B, S1A, and S1B), with local resolution reaching 2.79 Å in well-ordered regions (Figures 1C and 1D). The HA ectodomain is anchored to the viral membrane via a tripodal stalk region (Figure 1B).

**Figure 1.**
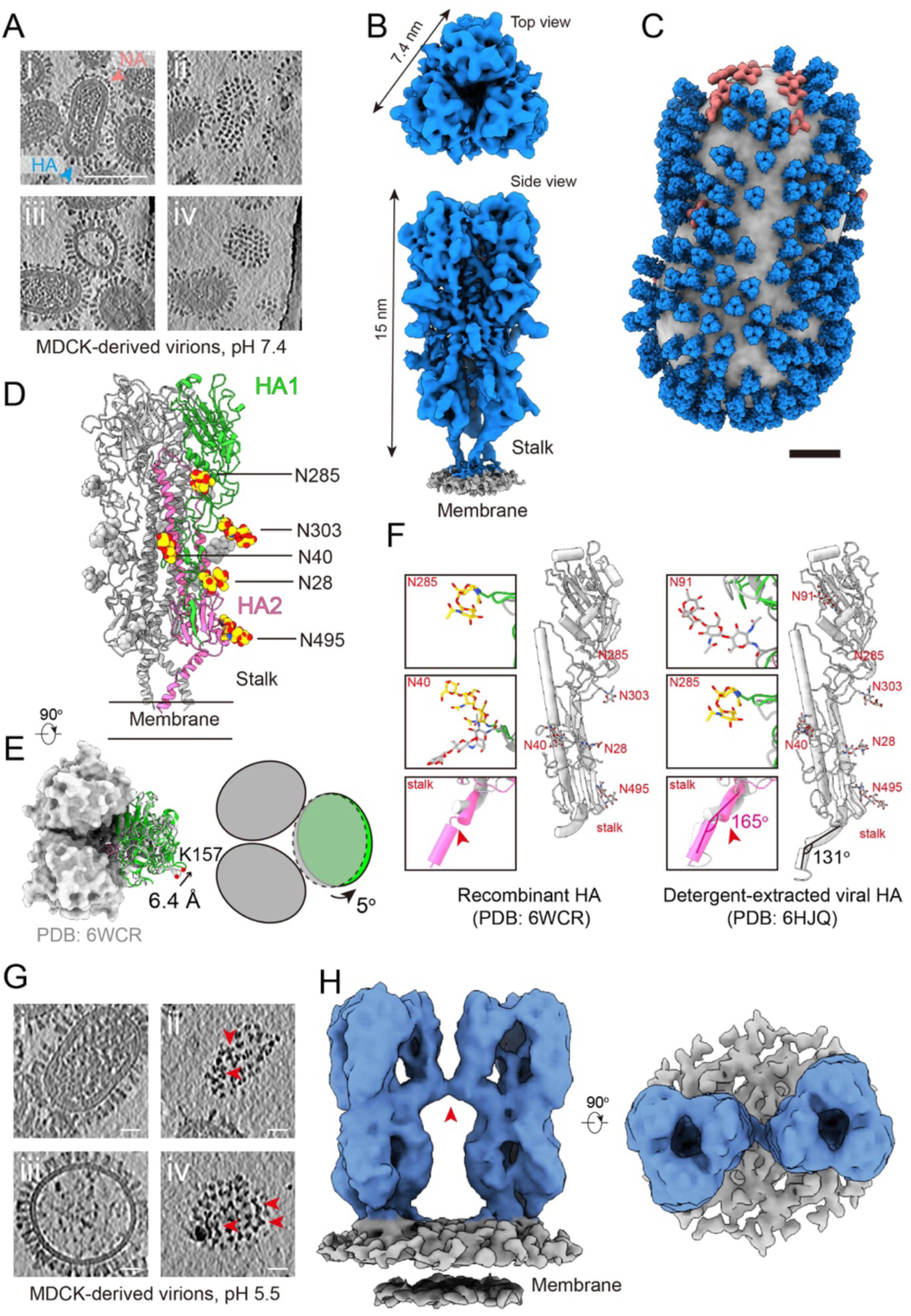
Structural analysis of HA on intact IAV virions derived from MDCK cells. **(A)** Tomographic slices (4 nm thickness) of virions at neutral pH, showing (i) An intact virion containing HA and NA on the viral envelope, a continuous M1 layer beneath the envelope, and densely packed RNPs; (ii) Top view of the virion in (i), where HA trimers are dense on the viral surface; (iii) Central section of a virion lacking the M1 while containing partial RNPs; (iv) Top view of the virion in (iii) where HA trimers are still dense despite the missing M1 and RNPs. Scale bars: 100 nm. **(B)** A 4.09 Å resolution on-virion structure of HA at low threshold, showing stalk densities. **(C)** A composite virion structure reconstructed by projecting the HA (blue) and NA (coral) structures onto their refined coordinates. The membrane is shown in gray. Scale bar: 20 nm. **(D)** An atomic model derived from the density map in (B). An HA monomer is colored in green (HA1) and pink (HA2). Spheres in red and yellow represent the five N-glycosylation sites visible on the map. Their identities from top to bottom: N285, N303, N40, N28, and N495. **(E)** Top view of the model, showing two protomers as surfaces in light grey, and the third as a cartoon. Compared to a model of the HA soluble ectodomain (PDB: 6WCR, dark grey), the on-virion HA1 model rotates outward by 5°, making the on-virion trimer slightly open. **(F)** Cartoon representations of a model of the HA soluble ectodomain (PDB: 6WCR, left), and a model of HA resolved from virions by detergent (6HJQ, right), shown as monomers. Their N-glycosylation sites are colored in red. The differences from our on-virion HA model are boxed, showing structural variations in the glycans (on-virion glycans in red and yellow) and the stalk region (on-virion stalks in pink). **(G)** Tomographic slices (4 nm thickness) of virions at acidic pH, showing (i) An intact virion; (ii) Top view of the virion in (i), where some HAs are paird (red arrowheads); (iii) Central section of a virion lacking the M1 and RNPs; (iv) Top view of the virion in (iii) where HA pairs are still visible. Scale bars: 20 nm. **(H)** Side and top views of a HA-dimer structure at pH 5.5. The HAs are connected by a densityon the HA1 region (red arrows).

To visualize glycoprotein organization on intact virions, we reconstructed an representative virion by projecting the refined HA and NA maps back onto their refined coordinates (Figure 1C). On PR8 virions, HA extends further from the viral envelope than NA, whereas the NA stalk region remain unresolved (Figure S2). NA molecules exhibited a tendency to cluster on the envelope, while HAs were densely distributed across the viral surface (Figures 1A and 1C). Notably, the structural features of NA differ from those reported for influenza strain X31^24^.

The 4.09 Å on-virion HA map enabled construction of a complete atomic model of the HA ectodomain (Figure 1D). The model resolves the stalk region and fifteen N-linked glycans, including those at N28, N40, N285, N303, and N495. Comparison with previously reported structures of recombinantly expressed PR8 HA ectodomain (PDB: 6WCR) and detergent-solubilized HA (PDB: 6HJQ)^25,26^ revealed three notable differences. First, alignment of HA2 subunits showed that the on-virion HA1 subunits adopt an outward, counterclockwise rotation of approximately 5°, corresponding to a 6.4 Å displacement at the trimer periphery, resulting in a more open HA head configuration (Figure 1E). Second, the N285 glycan is clearly resolved in the on-virion structure but absent from the soluble ectodomain model. Additionally, the N40 glycan projects away from the membrane in the on-virion model, in contrast to its orientation in soluble HA structures. Relative to the solubilized HA structure, HA1 glycosylation exhibited strain-specific variation, whereas HA2 glycosylation remained largely conserved. Third, the membrane-proximal stalk region of the on-virion HA comprises two α-helices connected by a short loop (residues 517-520), forming an inter-helix angle of approximately 165°. In contrast, the second helix is unresolved in recombinant HA, while the inter-helix angle is markedly kinked in the solubilized HA structure (131°) (Figure 1F). Collectively, these comparisons indicate that the on-virion HA structure captures an in situ conformational state characterized by HA1 “breathing”, a native HA2 stalk configuration, and more physiologically relevant glycosylation.

We next examined whether low-pH exposure promotes multi-HA assembly on intact virions. MDCK-derived virions were incubated at pH 5.5 (37 °C, 20 min) prior to cryo-ET imaging. The overall virion morphology and HA conformations closely resembled those observed under neutral conditions (Figure 1G). However, neighboring HAs frequently appeared as paired assemblies, independent of the presence of an underlying M1 layer (Figure 1G). STA of these particles resolved a dimer-of-trimers configuration connected by densities within the HA1 regions, hereafter referred to as the HA-dimer (Figure 1H).

### HA structures on egg-derived virions

To determine whether HA pairing is specific to virions propagated in cell culture, we next examined virions propagated in vivo. PR8 virions were amplified in embryonated chicken eggs and incubated under either neutral or acidic pH conditions for 15 min prior to cryo-ET imaging. Compared with MDCK-derived virions, egg-derived particles exhibited greater pleomorphism (Figures S3A and S3B), although virions from both sources displayed variability in internal structural integrity. In particular, viral particles lacking a continuous M1 matrix layer or containing incomplete ribonucleoprotein (RNP) assemblies were observed in both preparations (Figure S3A). Quantitative analysis revealed that 11% of MDCK-derived virions and 19% of egg-derived virions exhibited defects in M1 layer assembly (Figure S3C). Morphometric measurements showed that egg-derived virions were smaller (long, medium, and short axes: 113.95 ± 20.02 nm, 89.44 ± 7.15 nm, and 82.58 ± 7.12 nm) compared to MDCK-derived virions (124.15 ± 17.24 nm, 88.99 ± 6.58 nm, and 81.99 ± 7.43 nm) (Figure S3D).

Unexpectedly, locally ordered HA assemblies were frequently observed on egg-derived virions under neutral-pH conditions (Figure 2A). These oligomeric arrangements were detected irrespective of the presence of an underlying M1 layer and predominantly adopted pentameric or hexameric configurations, which further organized into extended lattice-like arrays. Examination of tomograms confirmed that these lattices were not induced by interactions with the air-water interface (Figure S4A). STA of approximately 94,000 HA particles extracted from 298 virions yielded a reconstruction at 3.58 Å resolution (Figures 2B, S5, S6A, and S6B). Notably, despite application of a mask encompassing only the central HA trimer during refinement, the reconstructed density remained connected to neighboring HA molecules at the HA1 periphery, indicating that the majority of HA trimers participated in oligomeric assemblies.

**Figure 2.**
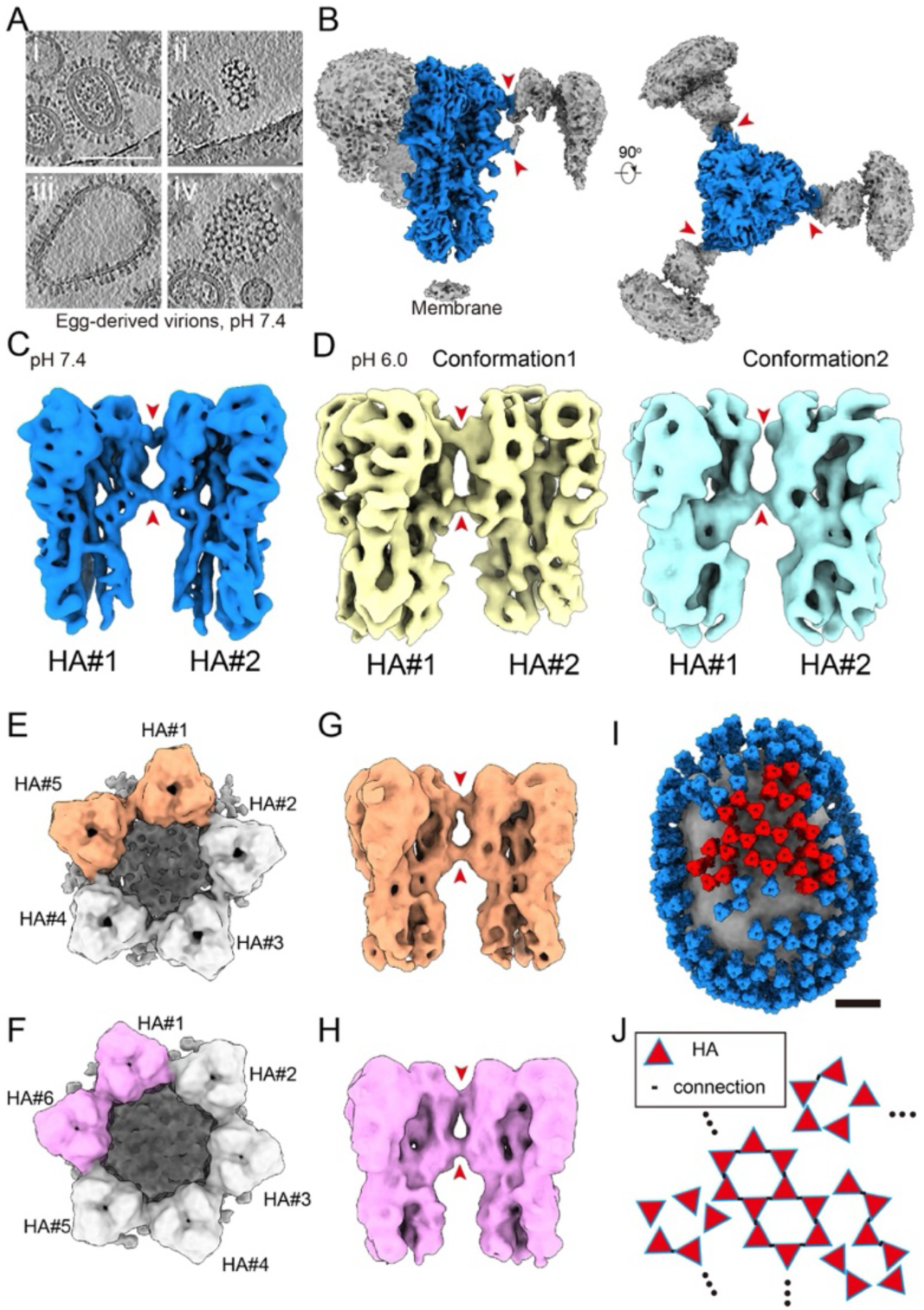
Structural analysis of HA on intact IAV virions derived from embryonated chicken eggs. **(A)** Tomographic slices (4 nm thickness) showing (i) a representative virion coated with HA and NA on the viral envelope, a continuous M1 layer beneath the envelope, and densely packed RNPs; (ii) Top view of a virion, where HA trimers arrange in lattice on the viral surface; (iii) Central section of a virion lacking the M1 and RNPs; (iv) Top view of the virion in (iii) where HA trimers still arrange in lattice despite the missing M1 and RNPs. Scale bars: 100 nm. **(B)** Side and top views of the HA structure determined from the viral surface. The central HA is colored blue, the adjacent HA and the membrane in grey. The central HA is connected to three neighboring HAs by two density bridges (red arrows). **(C-D)** Side view of a HA-dimer structure at neutral pH (C), and two HA-dimer structures at acidic pH (D). The HAs are connected by two densities on the HA1 region (red arrows). **(E-F)** Structures of an HA-pentamer and an HA-hexamer, with their composing HA-dimers (highlighted) displaying in side view **(G-H)**. **(I)** A composite virion structure reconstructed by projecting the HA structures onto their refined coordinates. A lattice composed of HA-pentamers and hexamers is highlighted (red). Scale bar: 20 nm. **(J)** A schematic illustrating that the lattice is built from the HA-HA connections.

An atomic model derived from the central HA density closely resembled the HA structure obtained from MDCK-derived virions (Figure S6C). Re-centering subtomograms to the midpoint between paired HAs enabled reconstruction of an HA-dimer map at 6.0 Å resolution (Figures 2C, S6D, and S6E). This map revealed two connecting densities between HA1 subunits, with the upper density appearing weaker than the lower. Under acidic conditions, egg-derived virions similarly exhibited HA-dimer assemblies (Figure S7A). Classification and refinement resolved two distinct HA-dimer conformations that differed primarily in the relative contact strength between HA1 subunits (Figures 2D and S7B).

To further characterize higher-order HA assemblies, subtomograms were re-centered on oligomeric clusters and subjected to classification. This analysis identified pentameric and hexameric assemblies of HAs. Initial reconstructions suggested structural flexibility within these oligomers, prompting additional classification to identify symmetrically ordered particles. Final reconstructions resolved ordered HA pentamers and HA hexamers at 8.0 Å and 9.7 Å resolution, respectively (Figures 2E, 2F, and S5). Dissecting the these higher-order oligomer structures revealed that they are constructed from HA-dimer building blocks (Figures 2G and 2H), with neighboring HAs engaging through interaction geometries similar to those observed in HA-dimers (Figures 2C and 2D). To illustrate how HA-pentamers and hexamers tile into flexible lattices on the viral surface through the HA interactions, we reconstructed an composite structure of an egg-derived virion with a patch of HA lattice highlighted (Figures 2I and 2J).

Collectively, these observations indicate that HA pairing represents a common organizational feature on egg-derived virions under both neutral and acidic conditions, while remaining relatively uncommon on MDCK-derived virions at neutral pH. Despite the diversity of oligomeric configurations, lateral interactions mediated by two HA1-HA1 interfaces constitute the fundamental structural basis of HA assembly. These interactions are likely weak and dynamic, enabling HA-dimers to adopt multiple geometries and accommodate the variable membrane curvature characteristic of pleomorphic IAV particles.

### Mechanisms of HA-dimer formation

To elucidate the structural basis of HA dimerization, we fitted the atomic model of the HA ectodomain into the 6.0 Å resolution HA-dimer map (Figure 3A). This analysis revealed two distinct interaction interfaces between neighboring HA1 subunits, hereafter referred to as interface 1 and interface 2. Interface 1 is located proximal to the receptor-binding domain (RBD) and involves residues 154-157, whereas interface 2 is positioned adjacent to the vestigial esterase (VE) domain and comprises residues 289-292 (Figure 3B). Both interfaces are primarily composed of polar and charged residues, including K57, H154, E155, K157, R162, H289, E290, and N292, suggesting that electrostatic and polar interactions could contribute to HA-dimer stabilization.

**Figure 3.**
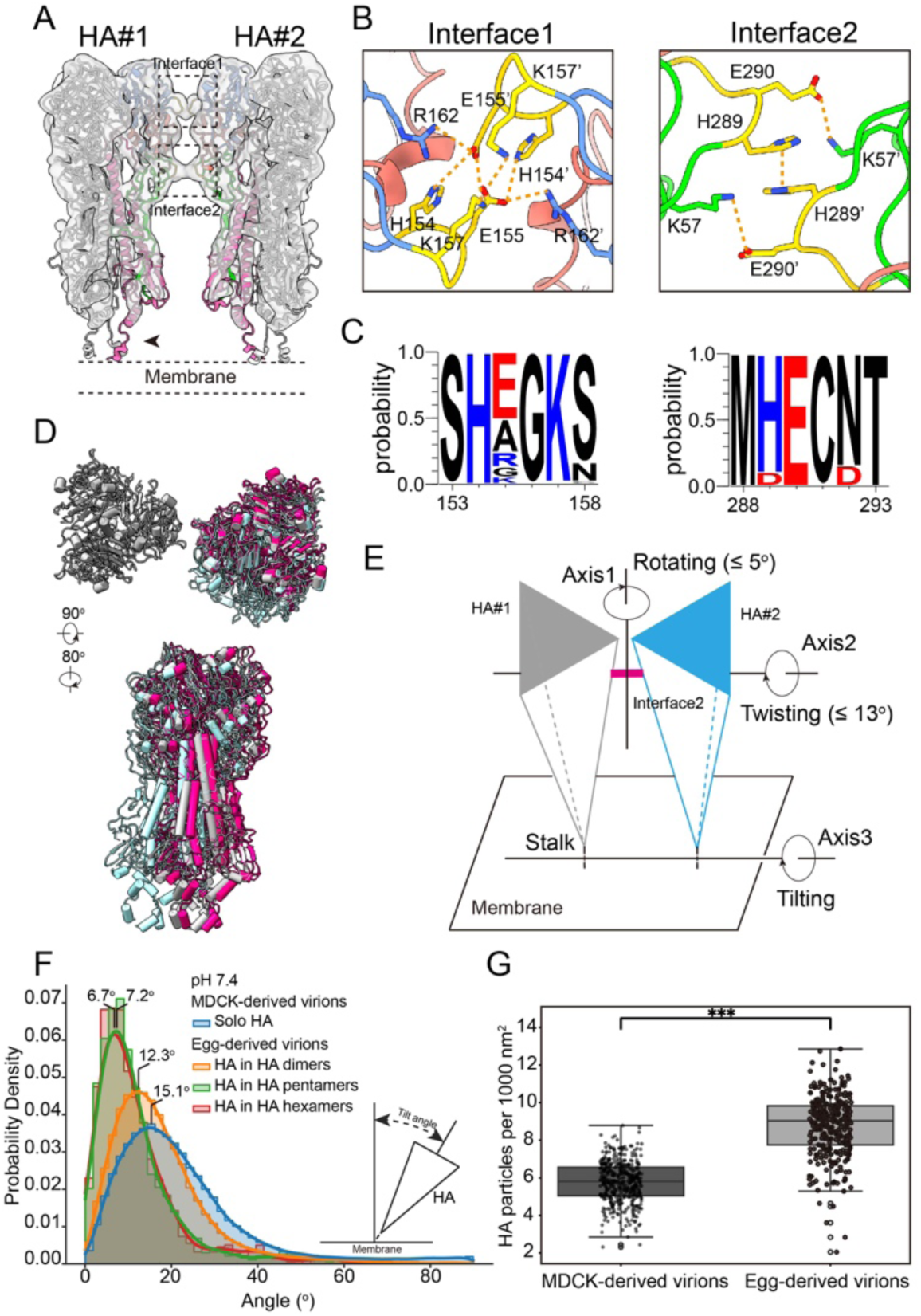
HA1 subunit mediates HA-pairing. **(A)** The on-virion HA model was fitted into the HA-dimer map from egg-derived virions at neutral pH. The HA protomer participating in HA-dimer formation is colored lime (HA1), pink (HA2), blue (RBD) and salmon (VE). Regions involved in Interface 1 and Interface 2 are highlighted with dashed boxes. **(B)** The zoomed-in view shows two loop segments (residues 154-157 and 289-292, gold) in close proximity at the HA-HA interface, suggesting potential contacts. **(C)** Sequence conservation analysis of amino acid residues potentially involved in forming Interface 1 and Interface 2, performed using WebLoGo. Charged residues are marked in blue (positively charged) and red (negatively charged). **(D)** The on-virion HA model was fitted into all conformations of HA-dimers. Representative colored models were aligned to HA#1 as reference (grey), illustrating the relative motion of HA#2 from top and side views. **(E)** A schematic illustrating three types of rotational motions of HA#2 (blue) relative to HA#1 (grey). For each rotation, an axis is drawn and the range of rotation is estimated from the fitting in (D). **(F)** Statistical distribution of HA tilting around axis3 in (E). The tilting angles of independent HAs, HAs from HA-dimer, pentamer and hexamers were separately analyzed. **(G)** Box plots of HA density (number per 1000 nm^2^ membrane area). Boxes indicate the interquartile range (IQR; 25th-75th percentiles) with the median as a horizontal line; whiskers span the 5th-95th percentiles; points denote outliers. Statistical significance was assessed using unpaired two-tailed t-tests; *** p < 0.001.

To assess the plausibility of these interactions, we measured inter-residue distances across the interfaces. At interface 1, the distances between E155 and opposing residues H154′, K157′, and R162′ were 3.8 Å, 5.3 Å, and 4.2 Å, respectively, with reciprocal spacing observed for E155′ relative to H154, K157, and R162. Given the effective range of electrostatic interactions, these geometries are consistent with moderate electrostatic stabilization. In contrast, interface 2 exhibited shorter inter-residue distances. The imidazole rings of opposing H289 residues were arranged in a nearly parallel orientation, with a center-to-center distance of 3.6 Å, indicative of potential π-π stacking interactions. Additionally, E290 residues were positioned to engage electrostatically with K57 (∼3.5 Å). Collectively, the tighter packing and complementary interaction geometries suggest that interface 2 provides stronger stabilization than interface 1.

To exclude the possibility that these interactions arise from passage-dependent mutations, we sequenced HA genes from both MDCK-derived and egg-derived virions. Sequence analysis confirmed that residues contributing to both interfaces were conserved across all samples (Figure S8). Broader conservation analysis across representative H1N1 strains further demonstrated that, although HA1 exhibits greater sequence variability than HA2, residues involved in HA1-HA1 interfaces remain relatively conserved and predominantly polar or charged (Figure 3C).

To characterize the conformational variability of HA-dimers, we fitted the HA ectodomain model into HA-dimer maps obtained under all experimental conditions and aligned the resulting models relative to a reference HA (HA#1) (Figure 3D). Notably, densities corresponding to interface 1 were variably resolved, whereas interface 2 densities remained consistently well defined (Figures 1H and 2D), further supporting a dominant stabilizing role for interface 2. The stronger interface 2 interactions permit limited relative motion between HAs within HA-dimers. Specifically, neighboring HAs exhibited rotational flexibility up to approximately ±13° about an axis passing through interface 2, and ±5° about an orthogonal axis (Figure 3E). These interfacial dynamics enable dynamic reshaping of HA-dimers. We next quantified HA orientation relative to the viral membrane. Analysis of the angle between each HA trimer’s threefold axis and the local membrane normal revealed considerable tilting flexibility. Scattered HAs exhibited tilting angles of up to ∼15°, whereas HAs engaged in HA-dimers, pentamers, or hexamers displayed progressively reduced tilting ranges (∼12°, ∼7°, and ∼7°, respectively) (Figure 3F). This reduction in orientational freedom indicates that oligomerization constrains HA mobility. The observed tilting behavior originates from structural flexibility within the membrane-proximal stalk region. In particular, the short linker (residues 517-520) connecting two α-helices in HA2 (Figure 1F) functions as a hinge that permits angular variation of the ectodomain. The combined effects of HA1 breathing, inter-trimer rotation, and stalk-mediated tilting establish a highly dynamic conformational landscape on the viral surface.

The six binding interfaces at the HA peripheries, together with HA’s structural plasticity, make HA multivalent in binding its neighbors. Formation of HA-dimers is likely spontaneous, as evidenced by their presence on isolated egg-derived virions at neutral pH. However, spontaneous lateral engagement of HAs requires close spatial proximity of neighboring trimers. Quantitative analysis revealed that egg-derived virions display a higher HA surface density than MDCK-derived virions. Egg-derived particles contained, on average, nine HA trimers per 1,000 nm² of membrane, with a mean center-to-center spacing of 76 Å. By comparison, MDCK-derived virions contained approximately six HA trimers per 1,000 nm², with a mean spacing of 87 Å (Figures 3G and S4B). These differences in molecular crowding likely contribute to the differential prevalence of HA oligomerization observed between virion populations.

### Disruption of HA1-HA1 interface impairs viral entry

To evaluate the functional significance of HA dimerization, we next investigated the contribution of the HA1-HA1 interaction interfaces to viral activity. Because structural analysis indicated that interface 2 constitutes the dominant stabilizing contact, we introduced alanine substitutions at all residues participating in this interface. The resulting mutant, designated HA_I2_3A (H289A, E290A, N292A), was designed to disrupt lateral HA interactions without perturbing global protein architecture (Figure 4A). To determine whether these mutations altered HA folding, we recombinantly expressed the HA_I2_3A ectodomain and determined its structure by single-particle cryo-EM. The reconstructed map revealed a conformation highly similar to that of our on-virion HA ectodomain, indicating that the interface mutations do not compromise HA structural integrity (Figures 4B and S9).

**Figure 4.**
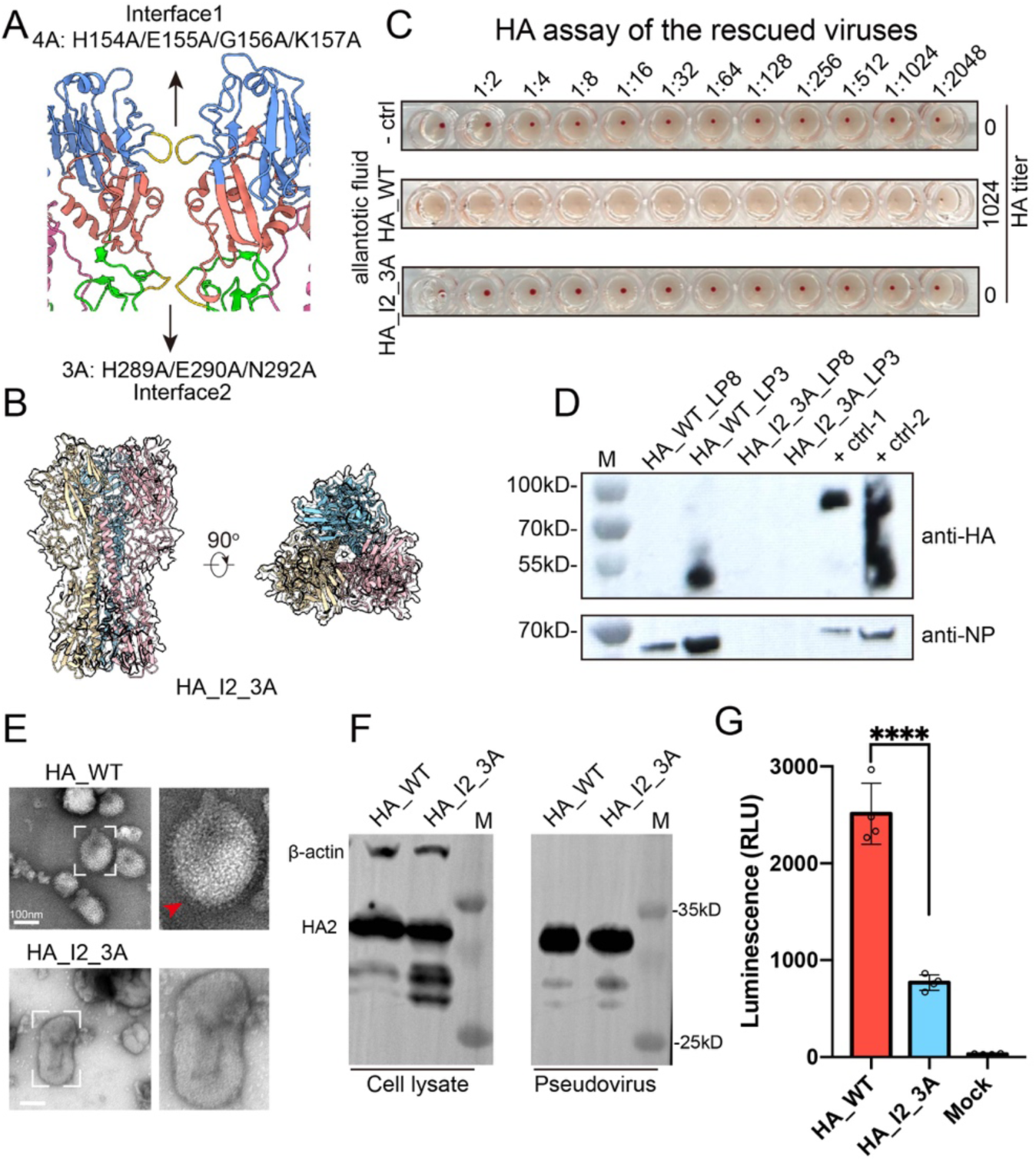
Disruption of the HA-HA interface reduces viral entry. **(A)** Mutated residues at HA1-HA1 interface. **(B)** Single particle reconstruction of the HA_I2_3A soluble ectodomain fitted with an HA soluble ectodomain model (PDB: 6WCR). The three protomers are colored yellow, blue, and magenta. **(C)** PR8 virus was rescued using an eight-plasmid reverse genetics system containing either HA_WT or HA_I2_3A. Chicken embryos were inoculated by the rescued virus and the harvested allantoic fluid was subjected to a hemagglutination assay. -Ctrl: Negative control using PBS. **(D)** Western blot analysis of HA and NP expression in the supernatant of HEK293T and MDCK co-cultures transfected with the eight-plasmid system. M: marker; LP8: transfection using Lipo8000; LP3: transfection using Lipo3000. +Ctrl: positive control using purified PR8 virus. **(E)** Chicken embryos were inoculated using the supernatant from (C), and the harvested allantoic fluid was purified for negative-staining EM. Particles highlighted by dashed boxes are enlarged on the right. Red arrows indicate PR8 virus particles. **(F)** Western blot analysis of HA in cells expressing HA_WT and HA_I2_3A (left) and in pseudoviruses harboring HA_WT and HA_I2_3A (right). **(G)** Infectivity characterization of pseudoviruses harboring HA_WT and HA_I2_3A. Data shown represent one of two independent replicate experiments, presented as mean values (bars) ± standard deviation (error bars) from n = 4 replicates (dots) in a single experiment. The significance of the difference was tested by t-test. ****, p < 0.0001. RLU, relative light unit.

We next assessed the impact of interface disruption on viral infectivity using an eight-plasmid reverse-genetics system to rescue recombinant PR8 viruses encoding either wild-type HA (HA_WT) or HA_I2_3A. Rescue of HA_WT virus resulted in robust viral recovery, as evidenced by strong hemagglutination activity, readily detectable HA and NP expression, and abundant virion production observed by negative-stain electron microscopy (Figures 4C-E). In contrast, no infectious virus was recovered from the HA_I2_3A construct. The failure of virus rescue indicates that residues comprising interface 2 play essential roles in the viral life cycle. However, because virus recovery depends on multiple replication steps, we sought to isolate the specific contribution of HA interactions to viral entry.

To this end, we generated HA-pseudotyped lentiviral particles harboring either HA_WT or HA_I2_3A. Immunoblotting and biochemical analyses confirmed comparable HA expression levels and equivalent pseudovirus production between constructs (Figure 4F). Pseudoviruses bearing HA_I2_3A exhibited markedly reduced luciferase reporter activity following infection of MDCK cells relative to HA_WT particles (Figure 4G). These results demonstrate that disruption of interface 2 specifically impairs HA-mediated viral entry, independent of effects on HA folding or particle production. Collectively, the data establish that lateral HA interactions are critical for efficient viral entry.

### HA-pairing facilitates viral fusion

We next examined whether lateral HA interactions directly contribute to membrane fusion efficiency. To this end, we performed quantitative cell-cell fusion assays using HEK293T cells co-expressing either wild-type HA (HA_WT) or interface mutants, together with TMPRSS2 to enable HA cleavage and activation (Figure 5A).

**Figure 5.**
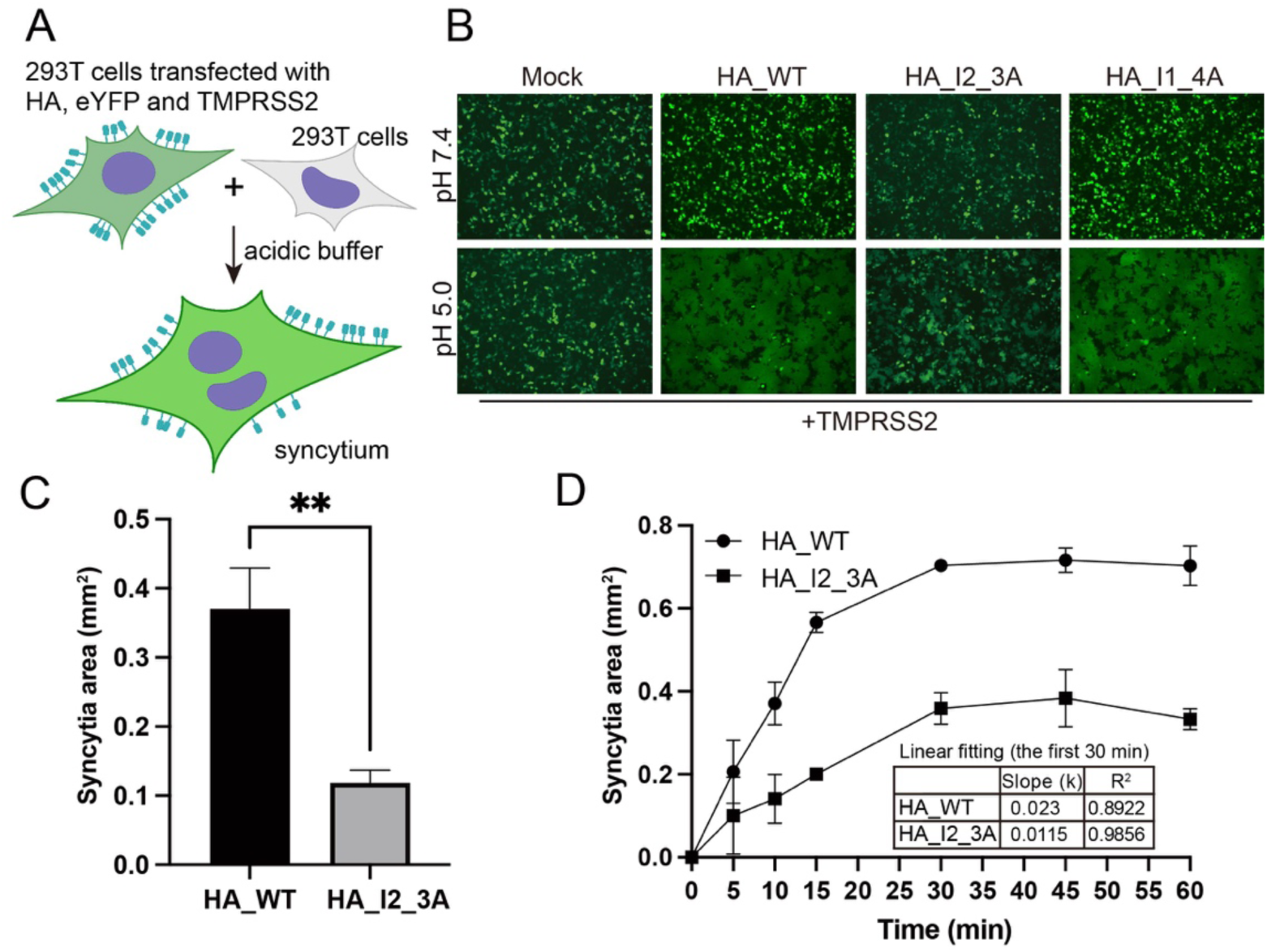
Mutations on the HA-dimer interface impair HA-mediated membrane fusion. **(A)** A schematic diagram of the cell-cell fusion assay. **(B)** Flourescence microscopy of HEK293T cells co-transfected with plasmids encoding either HA_WT, HA_I2_3A, or HA_I1_4A, together with human TMPRSS2 and eYFP. Mock: cells transfected with an equivalent amount of empty vector. **(C)** Quantification of syncytium areas (mm²) of HA_WT and HA_I2_3A transfected cells. Data are presented as mean values ± S.E.M from three independent experiments. Statistical analysis was performed using a t-test. **, p < 0.01. **(D)** Syncytium areas (mm²) of HA_WT and HA_I2_3A transfected cells were plotted over a series of time points with fitted curves. The points were linearly fitted for the first 30 minutes to calculate the slope.

Under neutral-pH conditions, no syncytium formation was observed for any construct, consistent with the requirement of low pH for HA activation. Upon exposure to fusion-permissive acidic conditions (pH 5.0), cells expressing HA_WT formed extensive syncytia, indicative of robust membrane fusion activity (Figure 5B). We first evaluated mutations targeting interface 1 (HA_I1_4A: H154A, E155A, G156A, K157A). Cells expressing HA_I1_4A exhibited fusion behavior comparable to HA_WT, suggesting that interface 1 contributes little to fusion efficiency under these conditions (Figures 5B and 5C). In contrast, disruption of interface 2 (HA_I2_3A) resulted in markedly reduced syncytium formation. Quantitative analysis of fusion areas revealed a substantial decrease in membrane fusion activity relative to HA_WT (Figures 5B and 5C). Time-resolved measurements further demonstrated that the initial rate of HA_I2_3A-mediated fusion was approximately twofold lower than that of HA_WT (Figure 5D). To exclude the possibility that reduced fusogenicity arose from altered protein expression or membrane trafficking, we performed immunofluorescence microscopy and immunoblotting analyses. These experiments confirmed that HA_WT and HA_I2_3A were expressed at comparable levels and efficiently localized to the plasma membrane (Figure S10). Together, these findings indicate that lateral HA interactions are not strictly required for membrane fusion but significantly enhance fusion kinetics. Disruption of interface 2 impairs the efficiency of HA-mediated membrane fusion, supporting a functional role for HA pairing in promoting cooperative fusion activity.

## Discussion

Our near-atomic resolution reconstructions of on-virion HA, together with structural characterization of HA oligomerization, distribution and mobility, reveal a highly dynamic organizational landscape on the IAV envelope. Compared with recombinantly expressed soluble HA ectodomains and detergent-solubilized HA structures, the in situ HA conformations determined here exhibit pronounced structural plasticity. In particular, HAs display a “breathing” motion characterized by outward dilation of HA1 subunits, as well as substantial tilting flexibility of the ectodomain relative to the viral membrane.

The observed HA tilting can be attributed to structural flexibility within the membrane-proximal stalk region, specifically the short linker connecting the two α-helices of HA2. This interpretation is consistent with previous structural studies of solubilized HA, which revealed multiple conformations differing in inter-helix angles^25^. Notably, our experimental findings are strikingly consistent with mesoscale molecular dynamics simulation of intact IAV particles, which similarly described HA breathing, tilting, and dynamic inter-spike interactions^27^. Together, these findings support a model in which HA structural plasticity represents an intrinsic feature of the native viral envelope.

Comparable spike dynamics have been reported for other enveloped viruses. Recent on-virion structures have revealed similar “breathing” conformations in SARS-CoV-2 S proteins^6,28^, ICV HEF^10^, and HSV-1 glycoprotein B^12^, as well as ectodomain tilting in SARS-CoV-2 S proteins^28,29^ and HIV-1 Env^30^. These observations suggest that structural flexibility may represent a general property of viral fusion glycoproteins. Importantly, such dynamics have direct implications for spike-spike interactions.

Our structural analyses indicate that HAs interact through multivalent, asymmetric interfaces. The presence of multiple peripheral interaction sites, comprising contacts of varying strengths, enables HA molecules to spontaneously assemble into dimers and higher-order oligomers. The predominance of HA oligomers on egg-derived virions further supports the idea that HA clustering represents a thermodynamically favorable process driven by molecular crowding and conformational flexibility.

In addition to intrinsic structural determinants, host-derived factors may modulate HA organization. Lipid raft microdomains, which are enriched in cholesterol and sphingolipids, have been shown to concentrate HA molecules during viral budding^31^. Receptor engagement has also been proposed to facilitate HA clustering, potentially via multivalent binding effects^11^. Indeed, HA-dimers observed in the presence of exogenous sialic acids closely resemble one of the HA-dimer conformations resolved here (Figure S11). However, the spontaneous formation of HA oligomers on isolated virions in the absence of supplemented receptors indicates that receptor binding is not essential for HA pairing. Instead, receptor interactions may stabilize or bias pre-existing HA assemblies. Antibody-mediated multivalent binding provides an additional mechanism capable of reshaping glycoprotein organization. Our recent cryo-ET studies have demonstrated that intact IgG molecules can cluster viral spikes through bivalent binding^29^, suggesting that immune factors may indirectly influence viral membrane architecture. Collectively, these considerations indicate that HA pairing arises from a combination of intrinsic structural plasticity and extrinsic environmental modulators.

Functional analyses presented here establish a direct mechanistic link between HA oligomerization and viral entry. Disruption of the dominant HA1-HA1 interaction interface abolishes virus rescue and markedly impairs HA-mediated entry and fusion kinetics. These findings support a model in which lateral HA interactions enhance the efficiency of membrane fusion rather than serving as an absolute requirement. We propose that HA pairing functions as a cooperative activation platform. By constraining spike mobility (Figure 3F) and promoting local spatial organization, HA oligomers may facilitate synchronized HA activation events, including HA1 dissociation, fusion peptide exposure, and membrane insertion^3,5^. Such coordinated transitions could reduce stochastic variability during fusion, thereby accelerating pore formation. This framework provides a mechanistic explanation for the reduced fusion kinetics observed upon interface disruption.

Beyond fusion energetics, HA clustering may influence receptor engagement, viral budding, and antigenic presentation. Because individual HA-sialic acid interactions are intrinsically weak, viral attachment relies on multivalent binding^32^. Locally ordered HA assemblies effectively generate clustered receptor-binding surfaces, thereby enhancing avidity and stabilizing virus-cell interactions^11,32^. HA clustering on infected cell membrane has also been found important for efficient IAV viral budding. HA accumulates in lipid raft microdomains that function as assembly platforms for virus morphogenesis and release. Mutating the HA2 TM domain to remove its association with lipid rafts resulted in virions showing reduced budding and greatly reduced infectivity, harboring less HA, and exhibiting decreased fusogenicity^31,33^. HA lateral interactions may also modulate antigenic presentation. The HA1-HA1 interfaces identified here are positioned adjacent to key antigenic sites, and oligomerization-induced tilting may alter epitope accessibility. Such structural rearrangements could influence antibody binding landscapes and immune recognition^34,35^.

Finally, comparison with oligomeric assemblies observed for other class I fusion proteins, including RSV F^9^, PFV Env^8^ and ICV HEF^10^ (Figure S12), highlights a potentially conserved structural principle: lateral contacts between membrane-distal head domains. Together, our findings reveal that viral fusion proteins are organized within a dynamic, cooperative network that shapes membrane fusion efficiency, receptor engagement, and antigenic architecture.

## STAR METHODS

### Virus propagation and purification

The laboratory strain influenza virus A/Puerto Rico/8/34 (H1N1) (PR8, ATCC VR95) were purchased from China Center for Type Culture Collection (CCTCC, Wuhan, China) and stored at −80 °C. Viruses were grown in 10-day-old specific-pathogen-free embryonated chicken eggs (Beijing Boehringer Ingelheim Vital Biotechnology Co., Ltd.) or Madin-Darby canine kidney (MDCK, ATCC CCL-34) cells for 72 h at 37 °C. Allantoic fluid or cell supernatant was harvested and subjected to low-speed centrifugation (2,000 g for 20 min at 4 °C) to remove cellular debris. Viruses were pelleted through a 33% (w/v) sucrose cushion by ultracentrifugation (112,000 g for 1.5 hour at 4 °C using a Beckman SW32 rotor). After aspiration of the supernatant, virus pellets were resuspended in phosphate-buffered saline (PBS, pH7.4). For further purification, a 10%-60% (w/v) sucrose gradient was applied. The virus band was extracted and dialyzed the sucrose, and a final concentration using centrifugal concentrators (Amicon, 0.5 mL volume, 100 kDa cutoff).

### Cryo-ET samples preparation

PR8 virions were mixed with 10 nm BSA-conjugated gold fiducial markers (Aurion, The Netherlands). A 4-µL aliquot of the mixture was applied to glow-discharged Quantifoil R2/2, 200-mesh holey carbon copper grids. Sample vitrification was performed in a Vitrobot Mark IV operating at 100% relative humidity and 8 °C. Grids were blotted for 4.5 s and plunge-frozen into liquid ethane, and subsequently transferred and stored in liquid nitrogen until data collection.

For samples subjected to acidic pH treatment, PR8 virions in pH 7.4 buffer (50 mM HEPES, 150 mM NaCl, 50 mM sodium citrate, pH 7.4) were mixed with 10 nm BSA-conjugated gold fiducial markers. The pH was adjusted to 5.5 or 6.0 using pH 3.0 buffer (50 mM HEPES, 150 mM NaCl, 50 mM sodium citrate, pH 3.0). The samples were incubated at 37 °C for 15 or 20 min to allow pH equilibration. Cryo-specimen preparation was performed as described above.

### Cryo-ET data collection

Cryo-ET data were acquired on a Titan Krios TEM (Thermo Fisher Scientific, Waltham, MA) operating at 300 kV and equipped with a Gatan K3 direct electron detector and a BioQuantum energy filter (slit width 20 eV). Tilt series were collected using a dose-symmetric scheme ranging from −60° to +60° with 3° increments, with a total cumulative dose of 131.2 e⁻/Å² and a defocus range of −2 to −4 µm. For PR8 virus samples derived from MDCK cells (neutral pH and pH 5.5 treated) and embryonated eggs (pH 6.0 treated), tilt series were acquired at a nominal magnification of 81,000× in super-resolution mode (calibrated pixel size, 0.5371 Å), except for MDCK-derived samples treated at pH 5.5, which were collected at 64,000× in super-resolution mode (calibrated pixel size, 0.68 Å). Automated data acquisition was performed using SerialEM^36^ with the PACEtomo script^37^. Tilt series for egg-derived PR8 virus samples at neutral pH were collected on a Titan Krios G4 microscope equipped with a Falcon 4i camera and a Selectris X energy filter (slit width 20 eV). Data acquisition was performed using the BIS-TOMO automated acquisition script developed by the National Multimode Trans-scale Biomedical Imaging Center based on SerialEM. Full data collection parameters are summarized in Table S1.

### Cryo-ET data processing

#### (1) Preprocessing

For PR8 virus samples derived from MDCK cells or embryonated eggs, tilt series data were preprocessed using FlyTomo^38^. Normal vectors to the membrane surface were generated using custom laboratory scripts employing Poisson surface reconstruction and sampled at half the nearest spatial distance of HA, defined as 4 nm (half of 8 nm). These normal vectors served as initial seeds for STA.

#### (2) STA of MDCK-derived PR8 virus

For STA of HA at pH 7.4, a total of 412,006 particles were extracted from the 8 × binned tomograms with a box size of 32³ voxels. For initial model generation, 15,091 particles were randomly selected from two representative tomograms. STA was conducted in Dynamo^39^, utilizing the PDB entry 1RU7 low-pass filtered to 20 Å as an initial template. The resultant average served as the initial model for a global Dynamo alignment encompassing all particles. After three iterative alignment rounds and the removal of overlapping particles, a dataset of 153,330 particles was obtained. Subsequently, this refined average, along with PDB entry 8E6J^40^ (also low-pass filtered to 20 Å), was employed as templates for multi-reference classification within Dynamo. This step facilitated the removal of aberrant or poorly aligning particles, yielding a final curated set of 107,446 particles. The refined particle set was exported to the Warp-RELION-M processing pipeline^41–43^. Tilt-series alignment parameters required by Warp were obtained from the FlyTomo outputs. Sub-tomogram reconstructions were computed in Warp, and three-dimensional (3D) classification and refinement were carried out in RELION 3 under C3 symmetry. Following removal of residual junk particles, 85,101 particles were carried forward into M. In M, multi-particle refinement of the tilt series and map refinement were performed over five sequential rounds, refining both geometric parameters (image and volume deformation) and per-tilt CTF parameters; 80 % of the available resolution range was employed during the first sub-iteration. Fourier shell correlation (FSC) between two independently refined half-datasets, local resolution estimation, and post-processing were conducted in M and RELION to assess map quality and resolution, yielding a final HA map at 4.09 Å resolution.

For STA of HA-dimers at pH 5.5, a total of 48 tomograms were reconstructed and subdivided into four independent Dynamo projects. Oversampled particles were extracted from bin4 tomograms using a box size of 32^3^ voxels, yielding an initial set of 333,012 particles. Multi-reference classification in Dynamo was first performed using coarse STA maps of HA and NA as references. After duplicate removal, particles were regrouped into four projects and subjected to a second round of multi-reference classification using coarse HA and HA-dimer templates. The HA-dimer template was generated by averaging 510 manually picked particles. Following an additional round of duplicate removal, 52,897 HA-dimer particles were retained. Particle coordinates were converted and imported into RELION 4 for multiple rounds of 3D classification and refinement, resulting in a final subset of 8,701 HA-dimer particles. Refined volumes were reconstructed at bin2 and subjected to C2-symmetric 3D refinement, yielding a final map at 13 Å resolution.

#### (3) STA of egg-derived PR8 virus

For STA of HA at pH 7.4, an initial set of 257,016 particles was extracted from 65 tomograms using an oversampling strategy in Dynamo. A robust initial reference was generated by aligning a random subset of 17,058 particles from five representative tomograms. This template was used to align the full dataset. Following alignment and distance-based duplicates removal, the dataset was reduced to 116,184 particles.

To select structurally homogeneous particles, multi-reference classification was performed within Dynamo, yielding a selected set of 94,330 subtomograms. These were exported to the Warp/RELION/M pipeline for high-resolution processing. Iterative 3D classification and refinement in RELION, followed by geometric refinement in M, resulted in a map of the HA at 3.58 Å resolution.

For the analysis of HA-dimer at pH 7.4, particles were manually re-centered and duplicates removed, expanding the dataset to 191,579 subtomograms. A subset of 28,427 particles representing coupled trimers was identified via 3D classification in RELION and subsequently refined in M, yielding a final map at 6.0 Å resolution.

For the analysis of HA-dimer at pH 6.0, a total of 297 tomograms were reconstructed and subdivided into 39 independent Dynamo projects. Oversampled particles were extracted from bin8 tomograms using a box size of 32^3^ voxels. Initial alignment was performed in Dynamo with C1 symmetry, using a coarse STA map of HA trimers as the reference. After duplicate removal, particles were regrouped into 13 projects for multi-reference classification using HA and NA templates. The HA-classified particles were realigned with C3 symmetry, applying a cylindrical mask. Following another round of duplicate removal, a total of 949,189 unique HA particles were retained.

To extract HA-dimers, we used a coarse STA map of the HA-dimer derived from MDCK-derived PR8 virions at pH 5.5, which exhibited clear dimeric features. This map was used as a reference for multi-reference classification of the full dataset, yielding an initial set of 508,942 candidate HA-dimer particles. Coordinates were converted and imported into RELION 4, and particles were evenly divided into seven jobs for batch processing. Bin4-level reconstructions were subjected to 3D classification (T = 0.1, without alignment) into 10 classes. Two conformational classes (conformation 1 and conformation 2) were selected for further refinement. Each subset was refined at bin4 resolution with imposed C2 symmetry, followed by additional 3D classification and refinement. Final particle counts were 28,900 for conformation 1 and 19,201 for conformation 2. Refined volumes were then reconstructed at bin2 and subjected to C2-symmetric 3D refinement. Per-particle frame alignment was performed at bin1, followed by a final round of bin2 3D refinement. The final reconstructions of conformation 1 and conformation 2 reached resolutions of 9.2 Å and 11.0 Å, respectively. Local resolution estimation was performed using RELION 4.

To characterize higher-order lattice organization at pH 7.4, the dataset of 191,579 particles underwent hierarchical 3D classification in RELION. Successive rounds of classification isolated a sub-population of 55,441 particles, which were further sorted into classes representing HA-pentamer and HA-hexamer. The HA-pentamer class (initially 7,642 particles) was cleaned to 4,398 particles, then underwent refinement in M with duplicate removal, resulting in a final set of 2,051 particles which were further refined in RELION. Similarly, the HA-hexamer class (initially 8,443 particles) was cleaned to 2,424 particles, refined in M with duplicate removal, yielding a final set of 1,225 particles for final refinement in RELION. These procedures yielded maps of the HA-pentamer and HA-hexamer at 8.0 Å and 9.7 Å resolution, respectively.

### Model building and refinement

Initial models of the full-length HA trimer were generated using AlphaFold3^44^ and docked into the density in ChimeraX^45^. For both HA structures, regions not resolved in the corresponding maps were manually removed from the predicted models in Coot^46^.

For the MDCK-derived HA, the membrane proximal stalk region was resolved at lower local resolution of approximately 6 Å and is predicted to comprise two alpha helices. These helices were flexibly fitted into the density using molecular dynamics flexible fitting (MDFF)^47^.

For both structures, N-linked glycans were manually built into well resolved glycosylation densities. The resulting ectodomain models were refined in real space using PHENIX^48^, followed by manual correction of steric clashes and side chain rotamers in Coot. Final model validation was performed in PHENIX. Model geometry and refinement statistics are summarized in Table S2. Local resolution filtered maps were used for visualization in ChimeraX. Structural alignments and comparisons were performed using the *matchmaker* command in ChimeraX.

### HA sequence comparison of egg- and MDCK-propagated PR8 viruses

Viral RNA was extracted from PR8 virions propagated in embryonated chicken eggs or MDCK cells using a commercial viral RNA extraction kit (Tiangen, Beijing, China), according to the manufacturer’s instructions. Reverse transcription was performed using an IAV universal primer (5′-AGCAAAAGCAGG-3′), which anneals to the conserved 3′ terminus of influenza A viral RNA segments, together with ABScript III Reverse Transcriptase (ABclonal, Wuhan, China) to generate complementary DNA (cDNA). The HA gene was subsequently amplified with gene-specific primers using Phanta Flash Super-Fidelity DNA Polymerase (Vazyme, Nanjing, China). PCR products were purified by agarose gel extraction and subjected to nanopore sequencing (CWbio, Beijing, China). HA sequences obtained from egg-and MDCK-propagated viruses were aligned and compared for sequence analysis.

Protein sequences were aligned using the ClustalW web server. Sequence conservation was visualized using WebLogo with the following protein sequences obtained from GenBank: ABO32981.1, ABO33006.1, ABN59434.1, AAA43171.1, AAP34322.1, AFO64857.1, BAK86315.1, ACA96508.1, ACD37430.1, ABU50586.1, ACU44318.1, ABW23340.1, AAP34323.1, AFO64813.1, ABI20826.1, ABD77807.1, ABD61735.1, ADT78908.1, ABD15258.1, ABD62843.1, ABD77675.1, ABO38351.1, ABD62781.1, ACV49556.1, ACF54598.1, ADW93963.1, AGV29214.1, APC60198.1, ANH71223.1, ATY75300.1, AGV28853.1, ACR26723.1, ACQ99821.1, and AAD17229.1.

### Expression and purification of recombinant HA_I2_3A ectodomain

To obtain the ectodomain of HA_I2_3A, the HA gene from the PR8 strain was synthesized and codon-optimized for expression in a human system. A truncated construct encoding residues 1-530, lacking the transmembrane and cytoplasmic domains, was cloned into the pCDNA3.1 vector. The C-terminus was fused with a T4 foldon trimerization motif followed by 8×His, FLAG, and Strep tags. Site-directed mutagenesis was performed to introduce H289A, E290A, and N292A substitutions, yielding the final expression construct.

The plasmid sequence was verified by Sanger sequencing and transfected into HEK293F cells using PEI MAX (Polysciences, Warrington, PA) at a DNA-to-PEI mass ratio of 1:3. After 72 hours, the culture supernatant was collected and subjected to Strep-Tactin affinity chromatography for initial purification. The eluted protein was further purified by size-exclusion chromatography (Superdex 200 Increase 10/300 GL, Cytiva, Wilmington, DE) and buffer-exchanged into TN buffer (20 mM Tris, 150 mM NaCl, pH 7.5).

### Cryo-EM sample preparation, data collection and processing

Purified HA_I2_3A ectodomain was concentrated to approximately 0.33 mg/ml in TN buffer. Three microliters of the protein solution were applied to glow-discharged Quantifoil R1.2/1.3 300-mesh copper grids (Quantifoil, Jena, Germany). The grids were incubated for 15 s at 4 °C and 100 % humidity, blotted for 3 s, and plunge-frozen in liquid ethane using a Vitrobot Mark IV. Cryo-EM data were collected on a Titan Krios TEM operating at 300 kV and equipped with a Gatan K3 direct electron detector and a BioQuantum energy filter (slit width 20 eV). Movies were recorded in super-resolution mode at a calibrated pixel size of 1.074 Å, with a total electron dose of ∼50 e⁻/Å² distributed over dose-fractionated frames. Data processing were carried out in cryoSPARC^49^, yielding a final reconstruction at 3.4 Å resolution.

### Immunofluorescence microscopy imaging

To generate a plasmid for high-yield antibody expression, the VH and Vκ genes of MEDI8852^50^ were synthesized (Tsingke Biotechnology, Beijing, China) and sequentially cloned into the pAbVec2.0 and pAbVec1.1 (kindly provided by Prof. Linqi Zhang at Tsinghua University), respectively. The plasmid sequence was verified by sequencing and transfected into HEK293F cells using PEI MAX (Polysciences, Warrington, PA) at a 1:3 DNA-to-PEI mass ratio. After 4 days of transfection, the supernatant containing MEDI8852 was harvested and clarified by centrifugation (2,000 × g, 20 minutes). The antibody was purified using a HiTrap Protein A HP column (Cytiva, Wilmington, DE), followed by gel filtration chromatography on a Superdex 200 Increase 10/300 GL column (Cytiva, Wilmington, DE).

HEK293T cells were seeded on glass-bottom dishes (Cellvis, Mountain View, CA) and cultured in DMEM for 24 hours. Cells were then transfected with either empty vector (pcDNA3.1), or full-length constructs of HA_WT or HA_I2_3A (H289A/E290A/N292A), using Lipo8000 (Beyotime, Beijing, China) according to the manufacturer’s instructions. Twenty-four hours post-transfection, live nuclei were stained by adding Hoechst 33342 (Beyotime, Beijing, China) directly to the culture medium and incubating at 37 °C for 15 minutes, followed by two washes with pre-warmed PBS. To label the plasma membrane, cells were incubated with 500 µL of working solution from the Cell Membrane Green Fluorescent Staining Kit (DiO, Beyotime, Cat# C1993S) for 10 minutes at 37 °C in the dark. Excess dye was removed, and cells were washed with PBS 2-3 times. Subsequently, cells were fixed in 4% paraformaldehyde (PFA) for 10 minutes at room temperature, followed by three rinses with PBS. For immunostaining, cells were blocked in PBST containing 1% bovine serum albumin (BSA) for 1 hour at room temperature. After washing, cells were incubated overnight at 4 °C with a recombinant human anti-HA monoclonal antibody (MEDI8852, 1 mg/mL, diluted 1:100 in PBST containing 1% BSA). Following incubation, cells were washed three times with PBST (3 minutes each, protected from light), and then incubated with Abcam Alexa Fluor 647-conjugated goat anti-human IgG Fc secondary antibody (diluted 1:500 in 1% BSA-PBST) for 1.5 hours at room temperature. After three additional PBST washes, fluorescent images were acquired using a Zeiss LSM980 confocal microscope equipped with Airyscan mode (excitation wavelengths: 405, 506, and 647 nm; objective: 63× oil immersion).

### Cell-cell fusion assay

To assess the role of HA1-HA1 interactions in mediating HA-driven membrane fusion, a quantitative cell-cell fusion assay was performed. HEK293T cells were co-transfected with plasmids encoding either full-length wild-type (HA_WT) or mutant HA_I2_3A, human TMPRSS2, and enhanced yellow fluorescent protein (eYFP), all cloned into pcDNA3.1 backbone. Transfections were carried out in 12-well plates seeded with ∼90% confluent cells using Lipofectamine 8000 (Beyotime, Beijing, China), following the manufacturer’s protocol. After 24 h incubation at 37 °C in 5% CO₂, cells were imaged using an EVOS fluorescence imaging system (Thermo Fisher Scientific, Waltham, MA) to confirm transfection efficiency.

To induce fusion, cells were incubated in a pH 5.0 fusion buffer (50 mM HEPES, 150 mM NaCl, 50 mM sodium citrate, pH 5.0) at 37 °C for 10 min. The buffer was then replaced with standard culture medium, and cells were returned to the incubator for 1 h to allow syncytium formation. Post-fusion imaging was performed under identical settings. Brightfield and eYFP fluorescence images were merged using ImageJ to evaluate the extent of syncytia formation, as indicated by eYFP redistribution. Quantification of fusion areas and statistical analysis were conducted using GraphPad Prism.

### Recombinant viruses rescued through reverse genetics

The eight plasmids (pDZ-PB1, pDZ-PB2, pDZ-PA, pDZ-NP, pDZ-NA, pDZ-M, pDZ-NS-GFP, pDZ-HA) required for influenza virus rescue, containing the genetic information of the A/Puerto Rico (Mountain Sinai)/8/1934 virus strain, as well as the polymerase I promoter and terminator sequences. These plasmids were amplified in *Escherichia coli* DH5α cells and purified using a plasmid purification kit (Mei5bio, Beijing, China) according to the manufacturer’s protocol. For transfection, HEK293T and MDCK cells were seeded at a 2:1 ratio into 6-well plates and cultured to approximately 90% confluence. The plasmids were co-transfected using Lipofectamine 3000 (Thermo Fisher Scientific, Waltham, MA), following the manufacturer’s instructions. After 24 hours, the medium was replaced with post-infection DMEM medium, supplemented with 0.1% Gibco fetal bovine serum (Thermo Fisher Scientific, Waltham, MA), 0.3% (w/v) bovine serum albumin (MP Biomedicals, Santa Ana, CA), 20 mM HEPES buffer, 1 µg/mL TPCK-treated trypsin (Aladdin, Shanghai, China), and 1% penicillin-streptomycin. The cells were then incubated for an additional 48 hours. The supernatant was collected, clarified, and subjected to ultracentrifugation over a 33% sucrose cushion to isolate P0 generation virus particles. The P0 virus was used to infect 10-day-old embryonated chicken eggs, and the allantoic fluid was collected 72 hours post-infection. The virus was then concentrated and purified by ultracentrifugation to yield infectious P1 generation virus. Throughout each stage, biochemical experimental techniques were employed to detect the influenza virus. These techniques included hemagglutination assay, western blot, and negative-stain EM.

### Western blot analysis

For influenza virus rescue identification, allantoic fluid containing the virus progeny (P1) was used for western blot analysis. NP and HA proteins were detected using mouse anti-NP antibody (1:2000 dilution, Cat. No. 11675-MM03T; Sino Biological, Beijing, China) and mouse anti-HA antibody (1:2000 dilution, Cat. No. 11684-MM03; Sino Biological, Beijing, China), respectively. The approach ensures accurate identification of the viral proteins in the recombinant sample.

To quantify HA protein expression in HEK293T cells transfected with HA_WT, HA_I2_3A, or an empty plasmid as a control, cells were harvested 24 hours post-transfection. The cells were lysed in ice-cold RIPA buffer and incubated on ice for 30 minutes. Following incubation, lysates were centrifuged at 12,000 × g for 10 minutes at 4 °C to remove cell debris, and the supernatant was collected. HA expression was detected using rabbit anti-hemagglutinin antibody (1:5000 dilution, Cat. No. 86001-RM01; Sino Biological, Beijing, China). Protein levels were normalized to β-actin, and the relative expression of HA was quantified by calculating the ratio of HA to actin band intensities using densitometric analysis.

### Hemagglutination assay

Hemagglutination titers were determined using a standard twofold serial dilution protocol in a 96-well V-bottom plate. Briefly, 50 µL of PBS was added to wells in columns 2-12. A total of 100 µL of virus suspension was added to each well in column 1, except for one well designated as the negative control, which received 100 µL of PBS. Twofold serial dilutions were performed by transferring 50 µL from column 1 to column 2, mixing thoroughly by pipetting, and continuing the serial dilution through column 12 with fresh pipette tips at each step. The final 50 µL from column 12 was discarded. Following dilution, 50 µL of 0.5% (v/v) chicken red blood cells (RBCs) (SenBeiJia Biological Technology, Nanjing, China) in PBS was added to each well. Plates were gently tapped to mix and incubated at room temperature for 30-60 minutes until RBCs in the negative control formed a distinct pellet at the bottom of the well. The hemagglutination titer was defined as the reciprocal of the highest virus dilution at which complete RBC agglutination was observed.

### Influenza pseudovirus packaging and entry assay

Influenza HA retroviral pseudoviruses (HA-pseudoviruses) carrying a luciferase reporter gene were produced by HEK293T cells. At the day before transfection, HEK293T cells were seeded to wells of a 12-well plate and incubated at 37 °C, 5% CO_2_ in an incubator for 16-18 h. The cells were then co-transfected with 1.6 ug pNL4-3-Luc-R-E- (MIAOLING Biology, Wuhan, China), 0.1 ug TMPRSS2-pcDNA3.1 and 0.6 ug WT/mutant HA expressing plasmids HA/3A-pcDNA3.1 using Lipofectamine 3000 (Invitrogen, Carlsbad, CA). At 20 h post-transfection, cells were fed fresh medium containing 100 units/mL of a commercial neuraminidase (New England Biolabs, Beverly, MA) to induce the release of HA-pseudovirus from the surface of the producer cells^51,52^. The cell culture supernatant containing virions were collected at 48 h post transfection, and clarified by low-speed centrifugation (1000× g, 10 min) to remove the cell debris.

For HA-pseudotyped virus entry assays, MDCK cells were seeded on 96-well plates and infected with 100 ul/well of the pseudoviruses for 48 h. The luciferase signal (relative luminescence units or RLU) was detected with Luciferase Reporter Gene Assay Kit (Yeasen Biotechnology, Shanghai, China) according to the manufactural instructions.

To analyze the expression of WT and 3A in HEK293T cells, cells were co-transfected with pNL4-3-Luc-R-E-, TMPRSS2-pcDNA3.1 and HA expressing plasmids as described above. After transfection, cells were lysed in RIPA lysis buffer (Beyotime, Beijing, China), and clarified by centrifugation at 12,000 g at 4 °C for 10 min. The supernatants were subjected to SDS-polyacrylamide gel (GenScript Biotech, Nanjing, China) and transferred onto PVDF membrane (Merck, Germany). The membrane was then blocked with 5% (w/v) nonfat milk (Sangon Biotech, Shanghai, China) in PBST (1×PBS pH 7.4 and 0.1% Tween-20). Subsequently the membrane was incubated with the 1:2000 diluted rabbit anti-Influenza A Virus Hemagglutinin monoclonal antibody (86001-RM01, Sino Biological, Beijing, China) and the 1:4000 diluted HRP-conjugated goat anti-rabbit secondary antibodies (SA00001-2, Proteintech, Rosemont, IL). Finally, the membrane was visualized using a chemiluminescence imaging system with the SuperSignal Chemiluminescent Substrate Kit (Thermo Fisher Scientific, Waltham, MA).

To compare HA expression in WT and mutant pseudoviruses, the transfected cell culture containing pseudoviruses were collected at 48 h post transfection, which was clarified by low-speed centrifugation (1000 g, 10 min). The supernatant was then layered onto a 20% (w/v) sucrose cushion and purified by ultracentrifugation (SW32.1 rotor, 150,000 g, 2 h, 4 °C) using an Optima XE-90 ultracentrifuge (Beckman Coulter, Brea, CA). The pellet was resuspended in PBS (pH 7.4) and was used for subsequent Western blotting. For HA protein detection, the same primary antibody as described previously was used. For p24 protein detection, a rabbit anti-HIV-1 Gag-p24 polyclonal antibody (11695-RP02, Sino Biological, Beijing, China) diluted at 1:2000 served as the primary antibody. The secondary antibody used and the membrane imaging producer are the same as described above.

## Date availability

Electron microscopy maps have been deposited in the Electron Microscopy Data Bank under accession codes EMD-74866 (HA from MDCK-derived virions at pH 7.4; atomic model deposited as PDB 9ZVA) and EMD-68353 (HA-dimer from MDCK-derived virions at pH 5.5), as well as EMD-68337 (HA from egg-derived virions at pH 7.4; atomic model deposited as PDB 22IH), EMD-74862, EMD-74863, and EMD-74864 (dimeric, pentameric, and hexameric assemblies of HA from egg-derived virions at pH 7.4), followed by EMD-68338 and EMD-68339 (two HA-dimer conformations from egg-derived virions at pH 6.0).

## ACKNOWLEDGEMENTS

This work was supported in part from National Natural Science Foundation of China (#32171195 and #32241031) and Tsinghua University Dushi Fund (#2023Z11DSZ001). We thank Dr. Jianlin Lei, Dr. Fan Yang and Dr. Xiaomin Li from the cryo-EM Facility, Technology Center for Protein Sciences, Tsinghua University, for their support on cryo-EM/ET data collection. We thank the computational facility support on the cluster of Bio-Computing Platform (Tsinghua University Branch of China National Center for Protein Sciences Beijing). We thank Dr. Xiaojun Huang and Dr. Yan Zeng from National Multimode Trans-scale Biomedical Imaging Center for their support on cryo-ET data collection. We thank the assistance of Bingyu Liu at the Imaging Core Facility, Technology Center for Protein Sciences, Tsinghua University.

## AUTHOR CONTRIBUTIONS

S.L. conceived and supervised the project. Y.C. prepared the influenza viruses, proteins, antibodies, and cryo-samples. Y.C. and H.L. performed the fusion assays and immunofluorescence microscopy. X.T. provided the plasmids for virus rescue. Y.C. and Zirui.Z. performed the virus rescue and HA sequencing. H.Z. conducted the pseudovirus-based entry assays. Y.C., Z.Z. and J.X. collected and processed the EM data. Y.C. and R.L. built the atomic models. Y.C., Z.Z., C.P. and Y.S. analyzed the structures. Y.C., Zirui.Z., Z.Z. and S.L. wrote the manuscript. All authors critically revised the manuscript.

