## Supplementary Information for "A Dynamic Oligomerization Network Coordinates Hemagglutinin-Mediated Membrane Fusion on Influenza Virions"

This file includes:

Figures S1-S12

Tables S1-S2

### Supplementary Figures

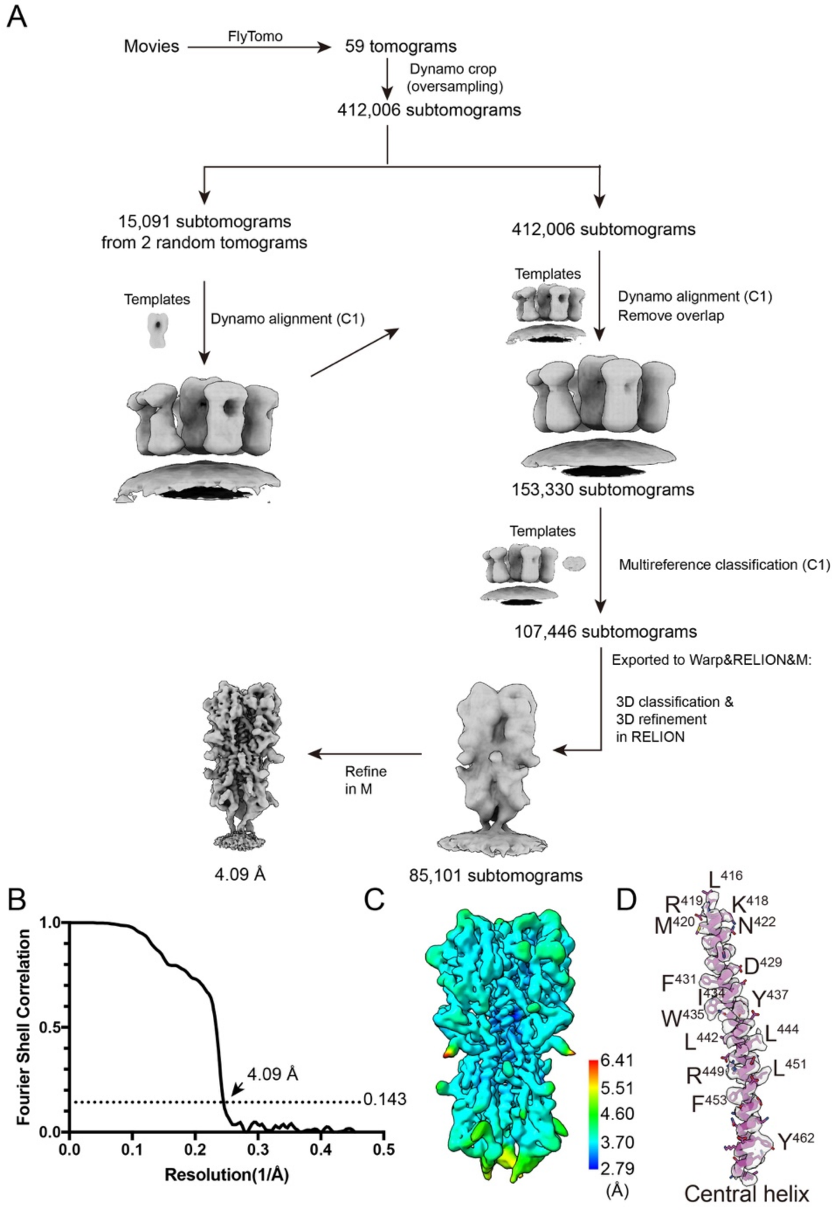


**Figure S1. Data processing workflow, FSC curves, and local resolution estimation of HA from MDCK-derived virions at neutral pH. (A)** Data processing workflow of HA timers. **(B)** Gold-standard Fourier shell correlation (FSC) curves for HA map. Dotted line indicated the 0.143 FSC cutoff. **(C)** Local resolution estimation using Relion and rendered by ChimeraX. **(D)** Density map of HA2 central helix was fitted the atomic model, demonstrating the representative side-chain densities.

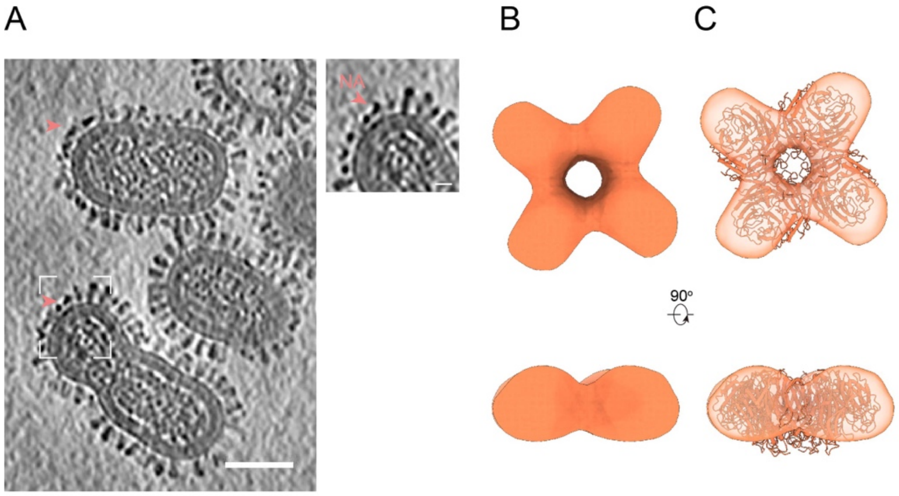


**Figure S2. Cryo-ET structure of influenza NA. (A)** Tomographic slice (4 nm thick) of representative MDCK-derived virions showing clustered NA on the viral membrane. Scale bar, 50 nm. The NA clusters highlighted by the dashed white box are zoomed in on the right. Coral arrowheads indicate NA molecules. Scale bar, 10 nm. **(B)** (upper) Top view and (lower) side view of the NA map. **(C)** (upper) Top view and (lower) side view of the N1 NA crystal structure (PDB: 8E6J) rigidly docked into the NA map.

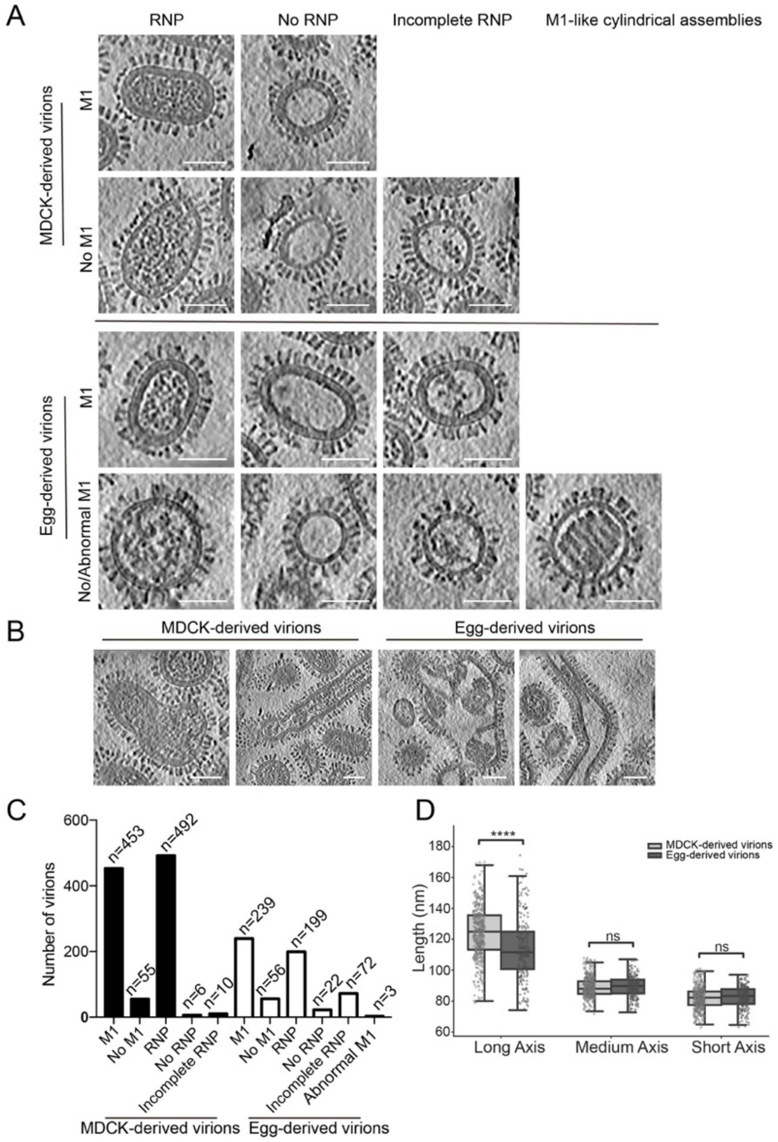


**Figure S3. Cryo-ET reveals structural heterogeneity of influenza A virions derived from MDCK cells and embryonated chicken eggs at neutral pH. (A)** Representative examples of pleomorphic IAV particles showing the presence, absence, or abnormal organization of the M1 layer, together with the corresponding internal organization of viral RNPs, including absent or incomplete arrangements. **(B)** Examples of irregular and filamentous virions. Scale bar, 50 nm. **(C)** Quantification of influenza virions with, without, or abnormal M1 layers and the corresponding RNP organization. **(D)** Box plot of the long, medium, and short axes of influenza virions derived from MDCK cells and embryonated chicken eggs. The top and bottom edges of the boxes represent the interquartile range (IQR), and the horizontal line within each box indicates the median. Whiskers span from the 5th to the 95th percentile, and individual dots represent outliers. Statistical significance was assessed using unpaired two-tailed t-tests. **** denotes $p < 0.0001$; ns indicates no statistically significant difference.

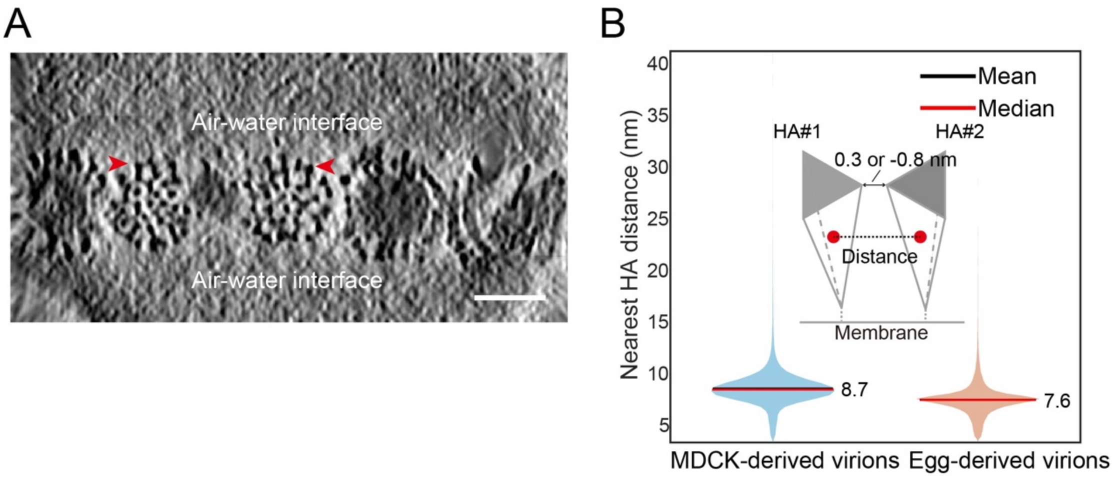


**Figure S4. Cryo-ET of HA on the surface of egg-derived influenza virions at neutral pH. (A)** A flipped representative tomogram slice (4 nm thick) showing the organization of HA trimers (red arrow) on the surface of egg-derived PR8 virions. Distinct HA oligomers are clearly visible away from the air-water interface. Scale bar: 50 nm. **(B)** The center-to-center distances between HA trimers and their nearest neighbors were calculated based on HA coordinates, and the statistical results are displayed as a violin plot.

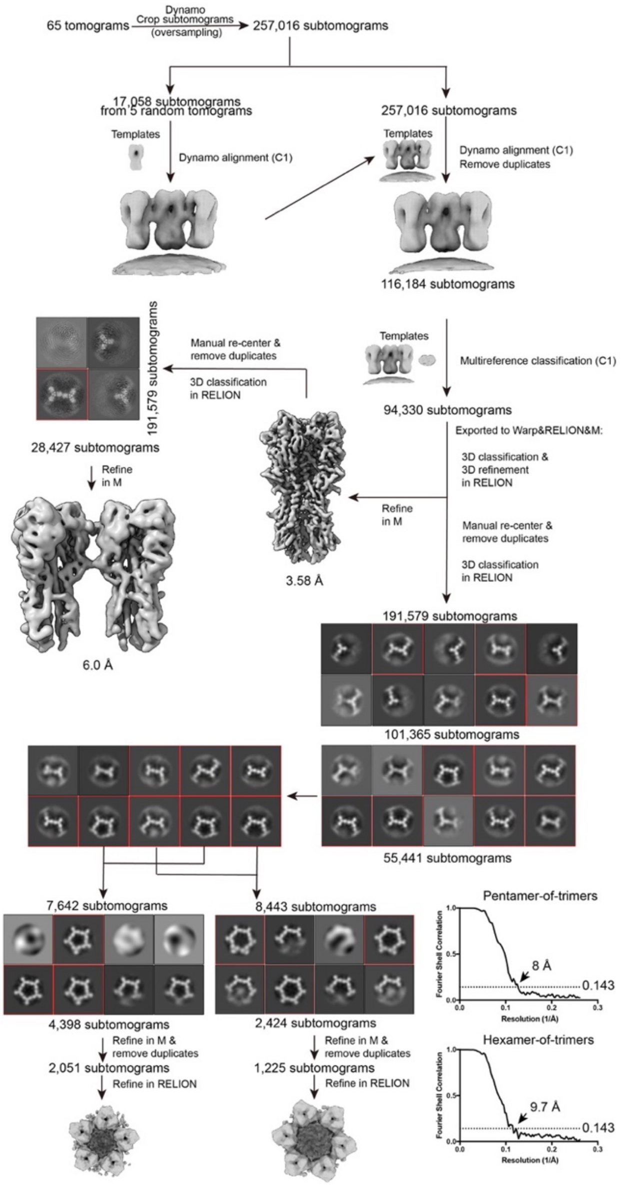


**Figure S5. Data processing workflow of HA, HA-dimer, pentamer, and hexamer from egg-derived virions at neutral pH.** Schematic overview of the data processing pipeline. 257,016 subtomograms were initially extracted. A subset of 17,058 particles (left branch) was used to generate templates for aligning the full dataset. After removing duplicates (116,184 particles) and Dynamo classification, 94,330 particles were retained.High-resolution refinement in Warp/RELION/M yielded a 3.58 Å map. A side branch identifies 28,427 particles corresponding to coupled trimers. To analyze oligomeric clustering, particles were re-centered and duplicates removed to form a dataset of 191,579 subtomograms. Hierarchical 3D classification isolated specific lattice geometries. A pentamer class was refined in M and RELION after duplicate removal, resulting in a final reconstruction from 2,051 particles. A hexamer class was similarly processed, resulting in a final reconstruction from 1,225 particles. Representative 2D class averages are displayed for the classification steps.

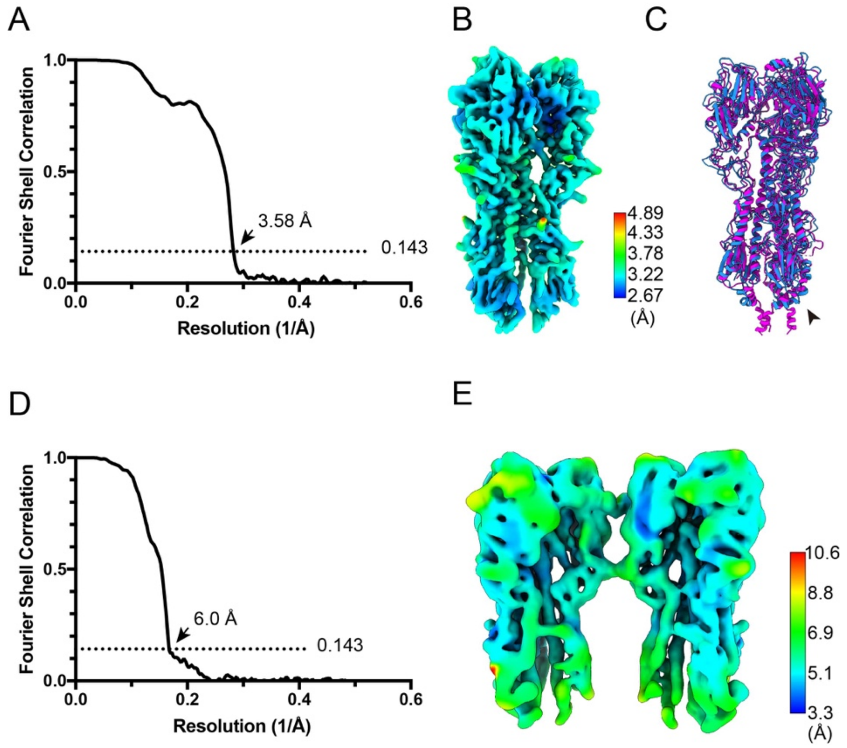


**Figure S6. FSC curves and local resolution of egg-derived on-virion HA and HA-dimer structures at neutral pH. (A, D)** Gold-standard FSC curves for the on-virion HA (A) and the HA-dimer (C). The dotted line marks the 0.143 FSC cutoff criterion. **(B**, **E**) Local resolution estimation of the on-virion HA (B) and the HA-dimer (D), performed in RELION and visualized with ChimeraX. **(C)** Cartoon representations of the on-virion HA atomic models derived from embryonated chicken eggs (blue) and MDCK cells (magenta). Black arrowheads indicate the differences in the stalk region.

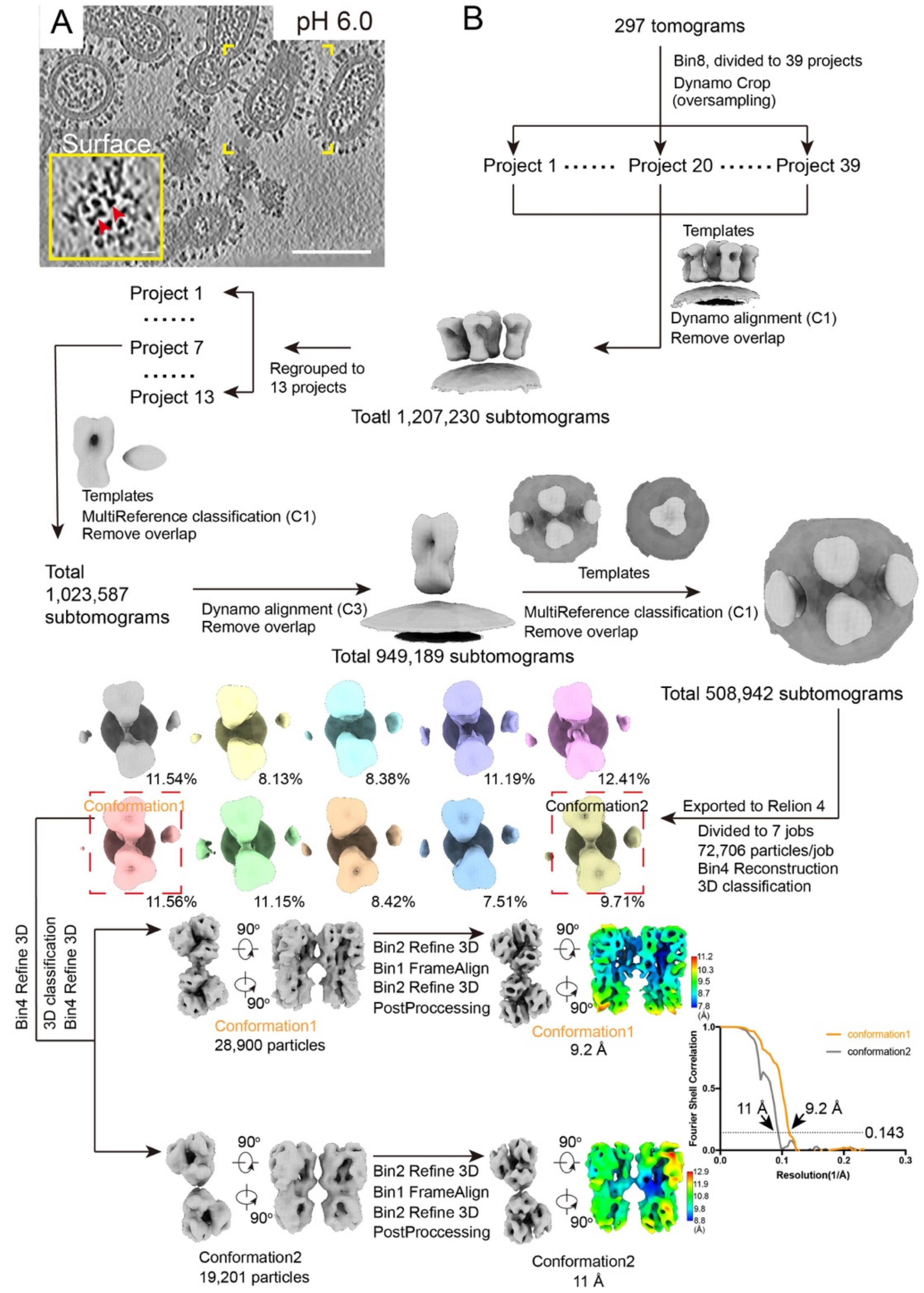


**Figure S7. Data processing workflow of HA-dimers from egg-derived virions at pH 6.0.** **(A)** A tomogram slice (thickness: 4 nm) showing the morphology of influenza virus and the distribution of surface glycoproteins at pH 6.0. The HA distribution on the viral surface within the yellow dashed box is zoomed in to the yellow inset at the bottom left. In the top view, HA trimers appear triangular, with adjacent HA trimers partially forming pairs through "vertex-to-vertex" interactions. Scale bars: 10 nm (inset); 100 nm (main image). **(B)** Data processing workflow of HA-dimers. Two major conformations were resolved at 9.2 Å and 11.0 Å resolution, respectively. Local resolution maps and FSC curves are shown.

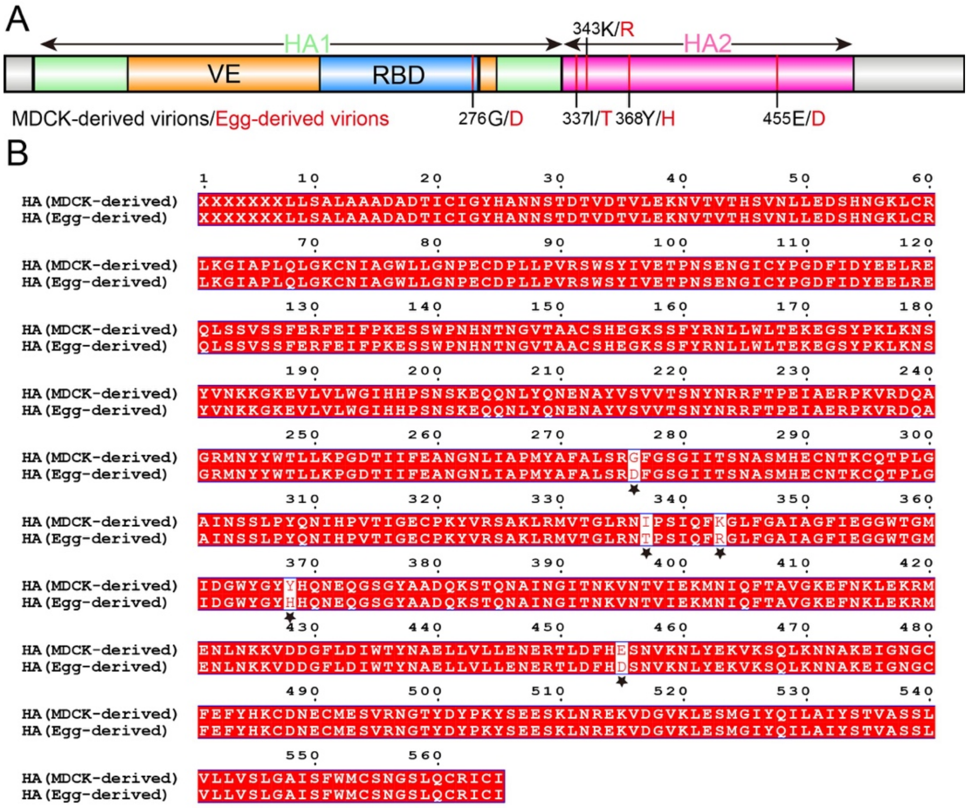


**Figure S8. Sequence comparison of HA from MDCK-derived and egg-derived PR8 virions.** **(A)** A schematic HA sequence generated using IBS 2.0, highlighting amino acid differences between MDCK- and egg-derived PR8 virions. **(B)** HA from MDCK-derived and egg-derived PR8 virions were obtained by reverse transcription and PCR amplification, followed by nanopore sequencing. The resulting nucleotide sequences were translated into amino acid sequences and aligned. Amino acid differences are indicated by black pentagrams.

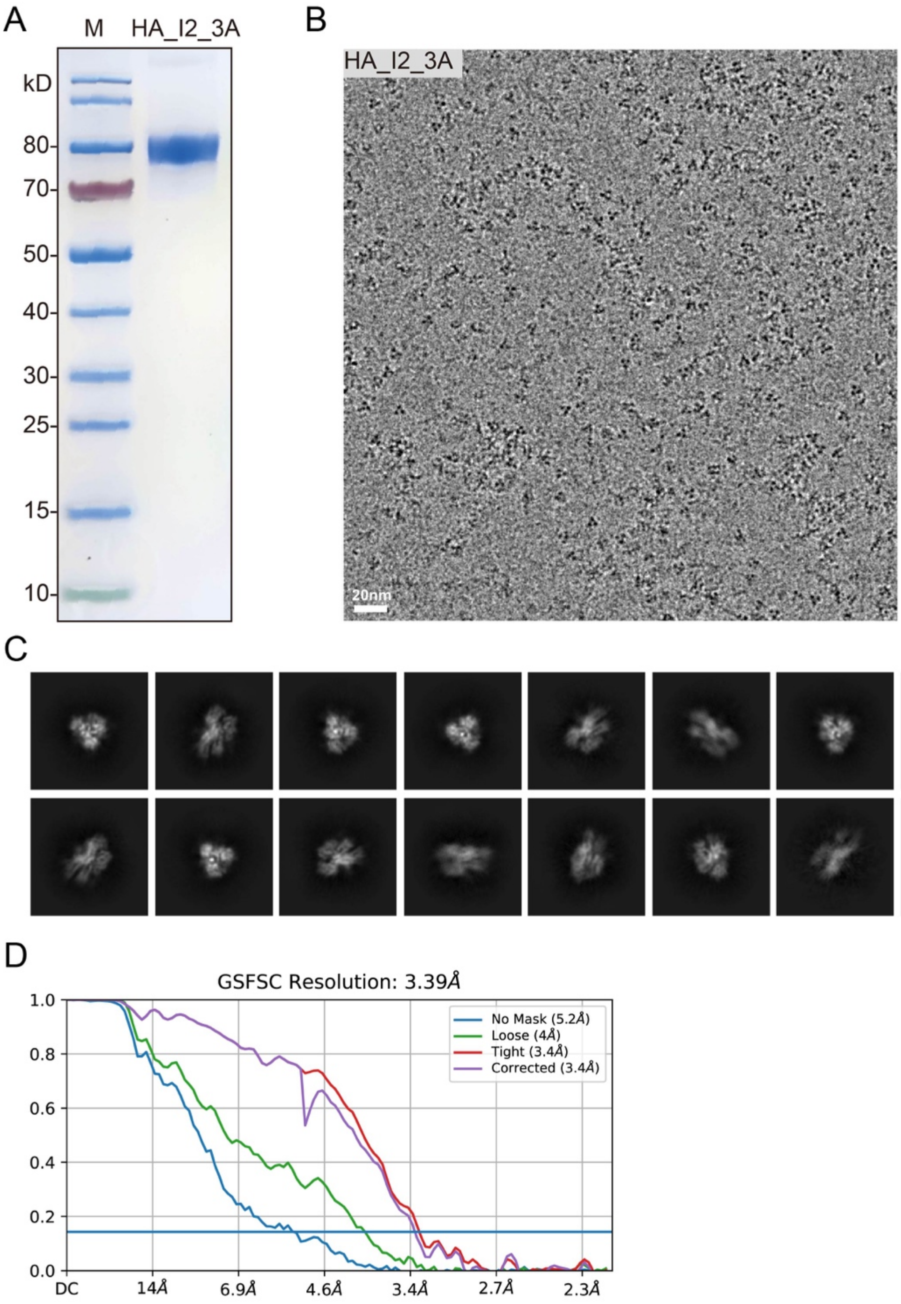


**Figure S9. Cryo-EM structure of HA_I2_3A. (A**) Purification of the HA_I2_3A ectodomain carrying the H289A/E290A/N292A triple mutation, recombinantly expressed in 293F cells. **(B)** Cryo-EM micrograph of HA_I2_3A particles. **(C)** Representative 2D class averages showing various particle orientations of HA_I2_3A. **(D)** FSC curves.

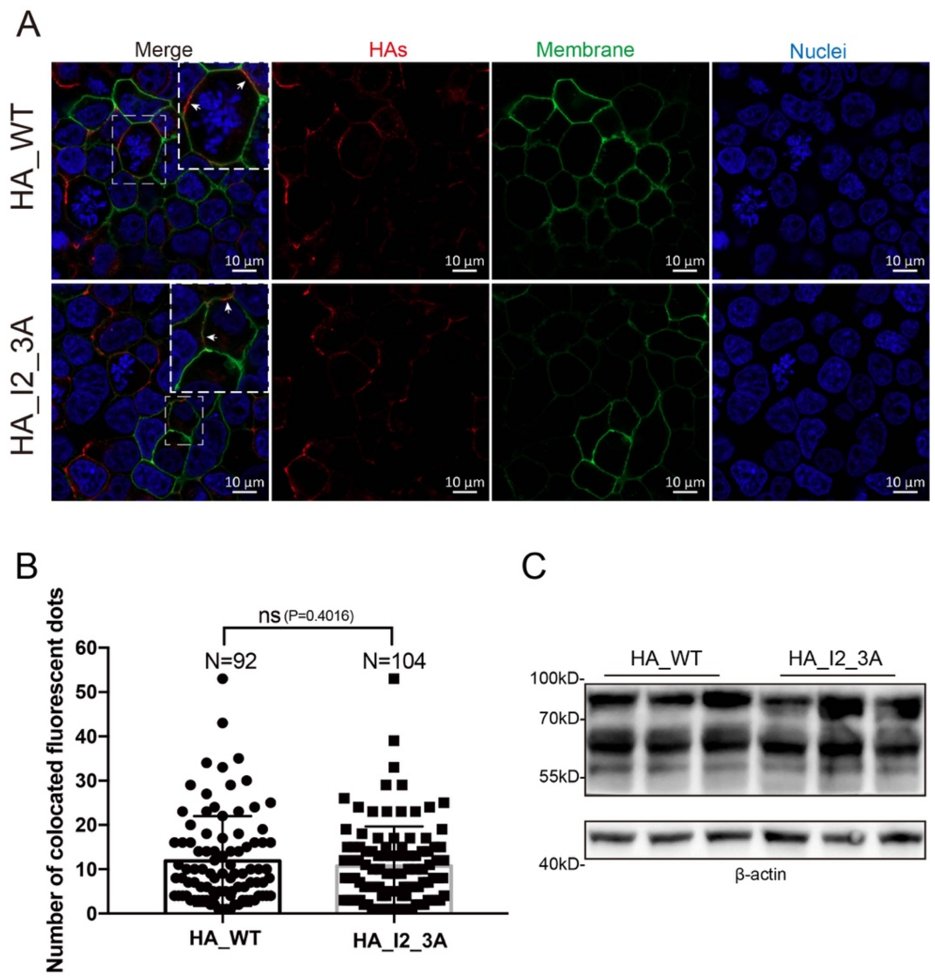


**Figure S10. Expression and quantification of HAs and mutants. (A)** Immunofluorescence co-localization analysis of HA_WT and HA_I2_3A with the cell membrane. HEK293T cells were fixed, and HA proteins along with their variants (red) were labeled by immunofluorescence staining. Nuclei were stained with Hoechst 33342 (blue), and the cell membrane was labeled with DiO (green). Both HA_WT and HA_I2_3A showed clear co-localization with the cell membrane. **(B)** The number of fluorescent signal points corresponding to HA_WT or HA_I2_3A that co-localized with the cell membrane was quantified using Imaris to assess their respective expression levels. Statistical analysis was performed using a t-test. **(C)** Western blot analysis of HA_WT and HA_I2_3A expression levels, with three replicates for each.

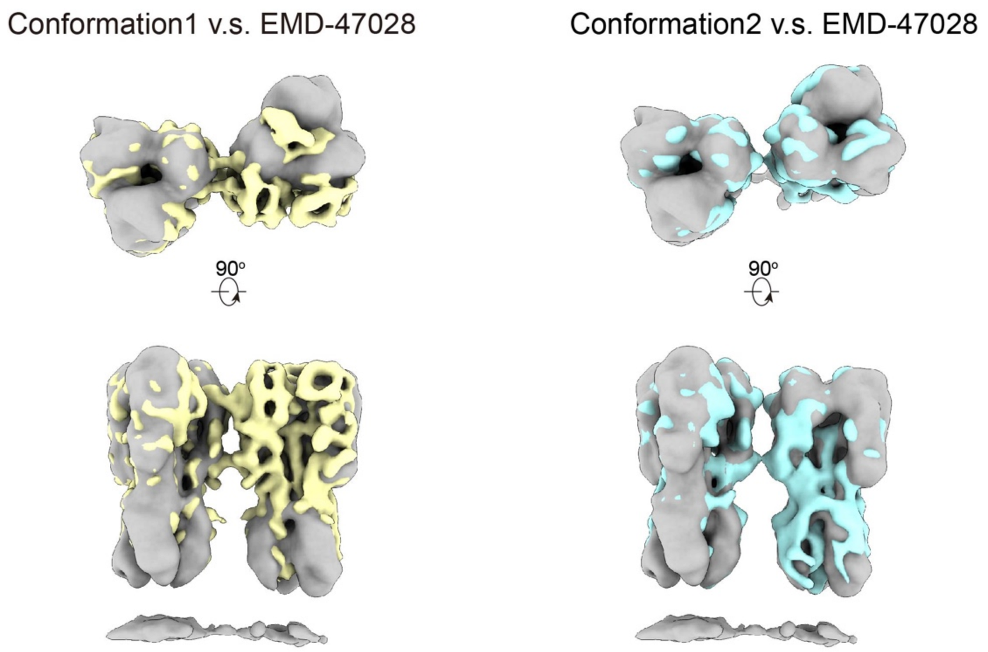


**Figure S11. Structural comparison of three conformations of HA-dimers.** Structural comparison of HA-dimer conformation 1 (yellow) and conformation 2 (cyan) on acid-treated egg-derived virions with the previously reported HA-dimer density (gray, EMD-47082) revealed distinct differences. When aligned by one HA trimer, the second HA in EMD-47082 exhibits a ~22 Å displacement at the globular head relative to conformation 1, pivoting around the interface 2 region. In contrast, the configuration of conformation 2 closely resembles that of EMD-47082, with minimal structural deviation.

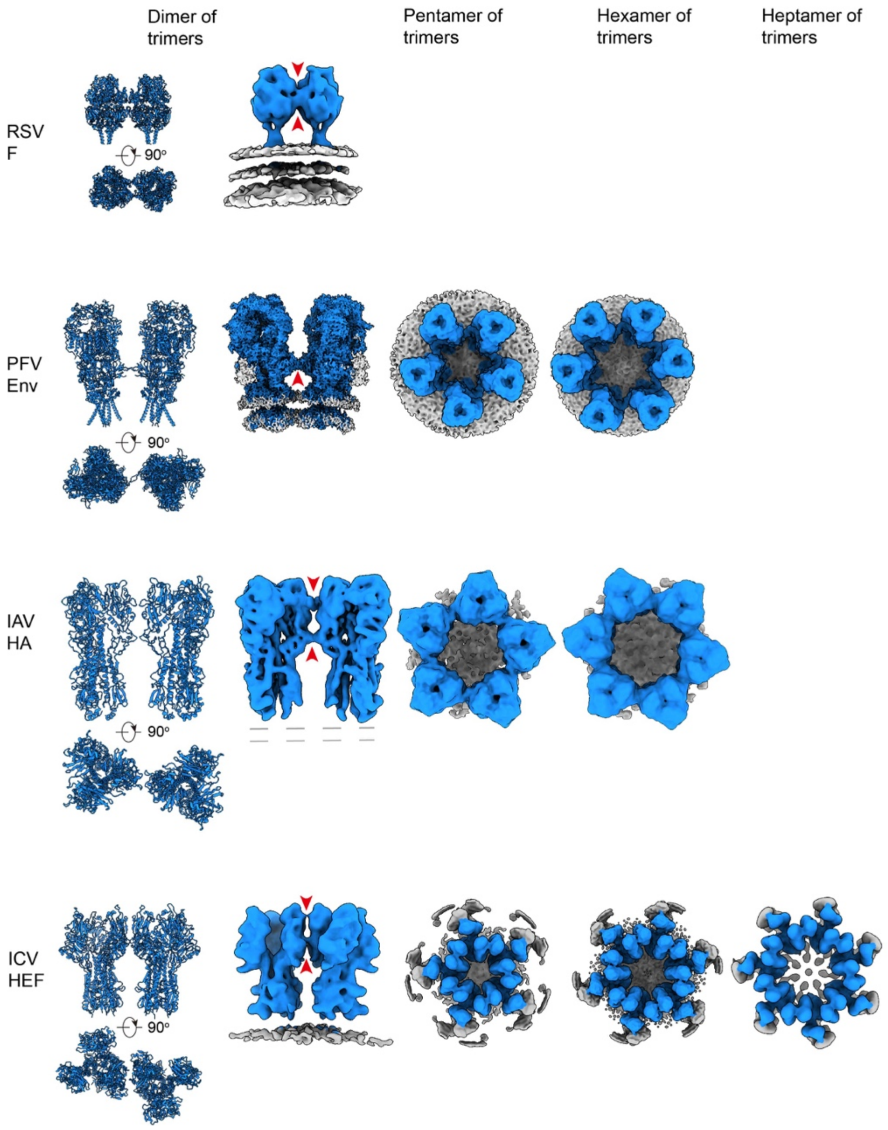


**Figure S12. Gallery of representative class-I viral fusion glycoproteins (GPs) from enveloped viruses in their prefusion conformations and higher-order oligomeric states.** Structures shown include RSV F (respiratory syncytial virus fusion glycoprotein; PDB 4JHW, EMD-44966), ICV HEF (influenza C virus hemagglutinin-esterase-fusion factor; PDB 6YI5, EMD-10810, EMD-17612, EMD-17001, EMD-17611), PFV Env (prototype foamy virus envelope protein; PDB 8OZJ, EMD-17311, EMD-17317, EMD-17318), and IAV HA (this study). Glycoproteins are depicted in blue, lipid bilayers in light gray. Red arrows highlight the inter-trimer contact sites between adjacent glycoproteins.

**Table S1. Cryo-ET data acquisition and reconstruction statistics of PR8 samples.**

| **Samples** | **MDCK-derived virions** | |
|---|---|---|
| pH | 7.4 | 5.5 |
| **Data acquisition** | | |
| Microscope | Titan Krios G3 | Titan Krios G3 |
| Voltage (keV) | 300 | 300 |
| Energy filter | GIF Quantum, Slit width 20 eV | GIF Quantum, Slit width 20 eV |
| Detector | Gatan Quantum K3 | Gatan Quantum K3 |
| Pixel size (Å) | 0.5371 (super resolution) | 0.68 (super resolution) |
| Defocus range (μm) | -2.0 to -4.0 | -2.0 to -4.0 |
| Tilt scheme | PACEtomo<br>Dose symmetric | Dose symmetric |
| Tilt range, step | $-60^{o}/60^{o}$, $3^{o}$ | $-60^{o}/60^{o}$, $3^{o}$ |
| Total dose ($e^{-}/Å^{2}$) | 131.2 | 131.2 |
| Frame number | 10 | 8 |
| Tomogram number | 59 | 48 |
| **Reconstruction** | | |
| | HA | HA dimer |
| Software | Dynamo 1.1.509, Warp, M, RLION3 | Dynamo 1.1.333, RLION4 |
| No. of subtomograms | 85,101 | 8,701 |
| Symmetry imposed | C3 | C2 |
| Resolution (Å) | 4.09 | 13.0 |
| FSC threshold | 0.143 | 0.143 |
| B-factor applied | -87 | -200 |

**Table S1 continued. Cryo-ET data acquisition and reconstruction statistics of PR8 samples.**

| **Samples** | **Egg-derived virions** | | | | | |
|---|---|---|---|---|---|---|
| pH | 7.4 | | | | 6.0 | |
| **Data acquisition** | | | | | | |
| Microscope | Titan Krios G4 | | | | Titan Krios G3 | |
| Voltage (keV) | 300 | | | | 300 | |
| Energy filter | Selectris X, Slit width 20 eV | | | | GIF Quantum, Slit width 20 eV | |
| Detector | Falcon 4i | | | | Gatan Quantum K3 | |
| Pixel size (Å) | 0.9576 | | | | 0.5371 (super resolution) | |
| Defocus range (μm) | -2.0 to -4.0 | | | | -2.0 to -4.0 | |
| Tilt scheme | BIS-TOMO<br>Dose symmetric | | | | PACEtomo<br>Dose symmetric | |
| Tilt range, step | -60°/60°, 3° | | | | -60°/60°, 3° | |
| Total dose ($e^-/Å^2$) | 131.2 | | | | 131.2 | |
| Frame number | 10 | | | | 10 | |
| Tomogram number | 65 | | | | 297 | |
| **Reconstruction** | | | | | | |
| | HA | HA dimer | HA pentamer | HA hexamer | HA dimer Comformation 1 | HA dimer Comformation 2 |
| Software | Dynamo 1.1.509, Warp, M, RLION3 | | | | Dynamo 1.1.333, RLION4 | |
| No. of subtomograms | 94,330 | 28,427 | 2,051 | 1,225 | 28,900 | 19,201 |
| Symmetry imposed | C3 | C2 | C5 | C6 | C2 | C2 |
| Resolution (Å) | 3.58 | 6.0 | 8.0 | 9.7 | 9.2 | 11 |
| FSC threshold | 0.143 | 0.143 | 0.143 | 0.143 | 0.143 | 0.143 |
| B-factor applied | -64 | -130 | -100 | -100 | -200 | -200 |

**Table S2. Validation statistics.**

| Refinement | HA (MDCK-derived virions) | HA (Egg-derived virions) |
|---|---|---|
| Initial model used (PDB code) | Alphafold | Alphafold |
| Model resolution (Å) | 4.4 | 3.8 |
| FSC threshold | 0.5 | 0.5 |
| Map resolution range (Å) | 2.79 to 6.41 | 2.67 to 4.89 |
| Map sharpening B factor ($Å^2$) | -87 | -64 |
| Model composition | | |
| Non-hydrogen atoms | 12615 | 11976 |
| Protein residues | 1524 | 1476 |
| B factor ($Å^2$) | | |
| Protein | 81.97 | 85.62 |
| R.m.s. deviations | | |
| Bond lengths (Å) | 0.002 | 0.005 |
| Bond angles (°) | 0.654 | 1.075 |
| Validation | | |
| MolProbity score | 2.18 | 1.92 |
| Clashscore | 15.03 | 10.85 |
| Poor rotamers (%) | 0.00 | 0.00 |
| Ramachandran plot | | |
| Favored (%) | 91.67 | 94.67 |
| Allowed (%) | 7.94 | 5.33 |
| Disallowed (%) | 0.40 | 0.00 |
